# Correcting PCR amplification errors in unique molecular identifiers to generate absolute numbers of sequencing molecules

**DOI:** 10.1101/2023.04.06.535911

**Authors:** Jianfeng Sun, Martin Philpott, Danson Loi, Shuang Li, Pablo Monteagudo-Mesas, Gabriela Hoffman, Jonathan Robson, Neelam Mehta, Vicki Gamble, Tom Brown, Tom Brown Sr, Stefan Canzar, Udo Oppermann, Adam P Cribbs

**Affiliations:** Botnar Research Centre, Nuffield Department of Orthopaedics, Rheumatology and Musculoskeletal Sciences, National Institute of Health Research Oxford Biomedical Research Unit (BRU), University of Oxford, Oxford, UK; Gene Center, Ludwig-Maximilians-Universität München, Munich, Germany; ATDBio Ltd (now part of Biotage), Oxford, UK; Chemistry Research Laboratory, Department of Chemistry, University of Oxford, Oxford, UK; Department of Computer Science and Engineering, Pennsylvania State University, University Park, PA, USA; Oxford Centre for Translational Myeloma Research University of Oxford, Oxford, UK

## Abstract

Unique Molecular Identifiers (UMIs) are random oligonucleotide sequences that remove PCR amplification biases. However, the impact that PCR associated sequencing errors have on the accuracy of generating absolute counts of RNA molecules is underappreciated. We show that PCR errors are the main source of inaccuracy in both bulk and single-cell sequencing data, and synthesizing UMIs using homotrimeric nucleotide blocks provides an error correcting solution, that allows absolute counting of sequenced molecules.

## Main

The inclusion of Unique Molecular Identifiers (UMI) in sequencing experiments creates a distinct identity for each input molecule, making it possible to correct sampling and PCR amplification bias. The use of UMI sequences pre-dates sequencing^1^, however including UMIs prior to library construction can improve the accuracy across almost all next-generation and third generation sequencing methods, including bulk RNA ^2, 3^, single-cell RNA ^4, 5^ and genomic DNA approaches^6, 7^. However, the accuracy of molecular quantification can be impacted by the varying sequencing quality of different platforms^8^. Moreover, different sequencing platforms require different PCR cycling conditions to generate adequate input material for sequencing, which can introduce UMI errors and lead to inaccurate molecule counts (Supplementary Fig. 1). Unlike sample barcodes for multiplexing or cell barcodes in single-cell sequencing, which can be whitelisted due to a limited pool of barcodes^9^, UMIs cannot be corrected using this approach as their synthesis is random. Therefore, UMIs are typically corrected using computational approaches^10^, concatemeric consensus sequencing^11^, or by bespoke UMI designs to aid error correction^12, 13^. Several computational approaches that leverage either hamming distances^14, 15^, graph networks^10, 12^, or thresholding on UMI frequency^4^ have previously been proposed to overcome PCR or sequencing errors within UMIs. However, none of these solutions have been experimentally validated and simulations suggest that UMI errors persist following computational demultiplexing (Supplementary Fig. 2).

We reasoned that utilising homotrimer nucleotides to synthesise UMIs would simplify error detection and correction by using a “majority vote” method (Fig. 1a and Supplementary Fig. 3). To enable more accurate error identification and better tolerance to indels, we developed a strategy that involves labelling each RNA molecule with an oligonucleotide containing a homotrimeric UMI located at the 5’ and/or the 3’ end, followed by sequencing using Oxford Nanopore Technologies (ONT), PacBio or Illumina sequencing platforms (Fig. 1a-d). UMIs are identified and errors are detected across the entire UMI sequence by comparing trimer complementarity. To correct these errors, we use a “majority vote” method where the most common nucleotide within the trimer is selected (Fig. 1e). Our simulations demonstrate that using trimers alone significantly improves the accuracy compared to relying solely on computational demultiplexing methods (Supplementary Fig. 4). To further improve the homotrimeric majority vote error removal, we utilised a combinatorial optimisation approach (Supplementary Fig. 5). To accurately identify translocated reads and distinguish them from chimeric artefacts, we use dual UMIs attached to both ends of the cDNA. This method aids in computationally removing chimeric artefacts (Fig. 1f and supplementary Fig. 6). By implementing our homotrimeric correction approach, we were able to improve the error detection and recover simulated UMIs that match the ground truth (Fig 1g, Supplementary Fig. 7)^12^.

**Figure 1:**
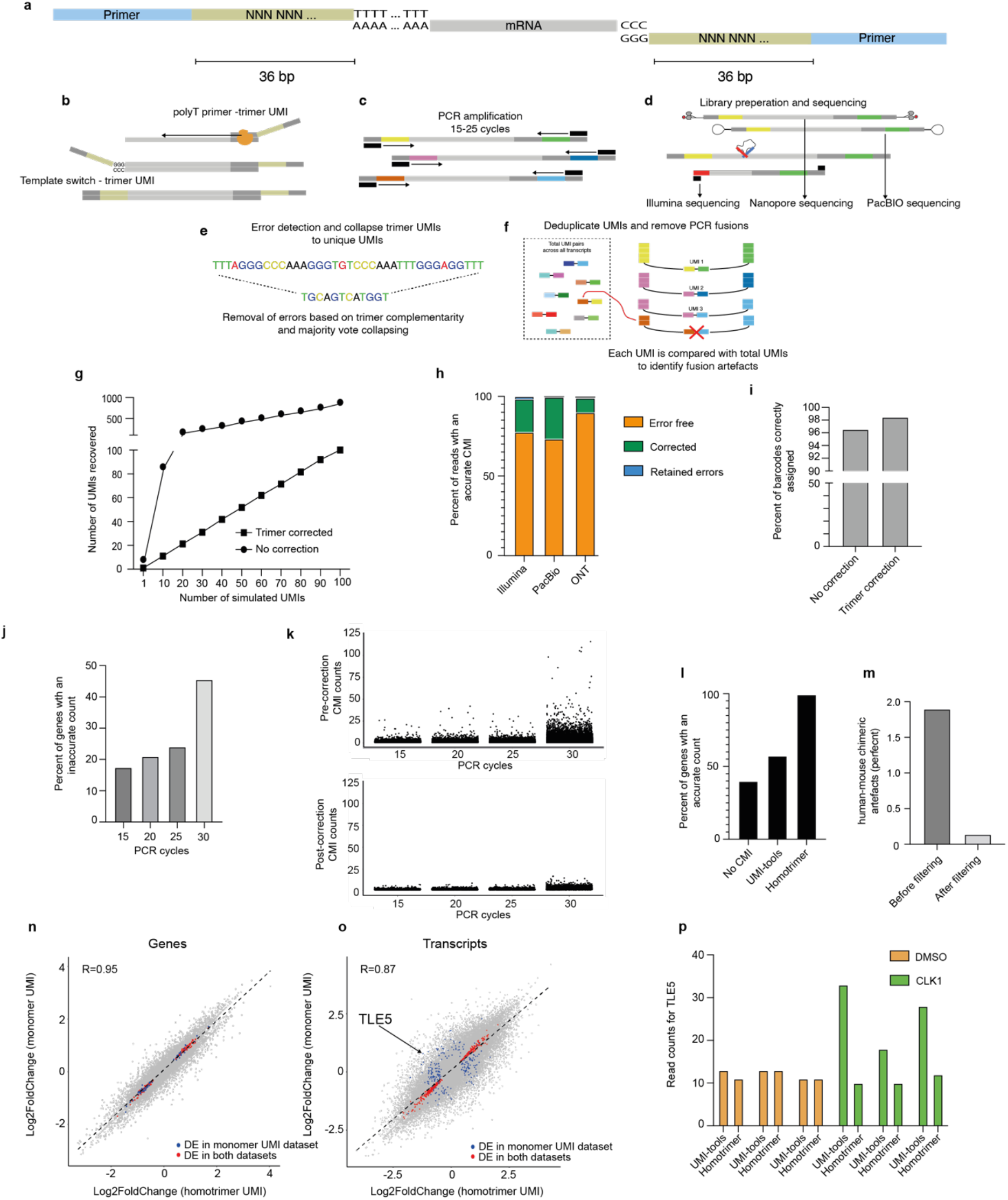
**a**, Schematic showing attachment of 3’ and 5’ UMIs to mRNA. **b**, The 3’ end of the mRNA is captured using an polyT oligo containing one homotrimer UMI of length 12, and optionally the 5’ end UMI is attached during template switching. **c**, PCR amplification is performed. **d**, Illumina, ONT or PacBio library preparation is then performed. **e**, Errors are then corrected using the homotrimer correction method. **f**, Removal of chimeric artefacts are performed using dual 3’ and 5’ UMIs. **g**, Simulated data showing the effect of deduplicating UMIs following homotrimer correction. **h**, Percent of CMIs that are correctly sequenced and then error corrected using homotrimer correction across Illumina, PacBio and ONT sequencing platforms. **i**, Barcode assignment using homotrimer barcodes before and after majority vote correction. **j**, Percent of genes with an accurate CMI count following increased PCR cycles of the same sequencing library. **k**, CMI counts plotted for each transcript showing the numbers of counts per transcript (the ground truth count for each transcript should be equal to 1). **l**, Percent of genes with an accurate CMI count following counting using UMI-tools correction and homotrimer error correction. **m**, The number of artefactual human-mouse chimeric reads removed using the dual UMI error correction approach (mcl-UMI). **n-p**, RM82 sarcoma cells were treated with DMSO or SGC-CLK-1 for 24 hours. Scatter plot of the log2 fold changes obtained from randomly collapsing each sequenced trimer UMI and then applying UMI-tools deduplication versus the log2 fold changes obtained from homotrimer UMI correction and counting for genes (**n**) and transcripts (**o**). Red points indicate the overlapping significant genes/transcripts and blue points indicate genes/transcripts that were disconcordantly significantly differentially expressed. **p**, TLE5 read counts showing the expression for DMSO and SGC-CLK1 following the application of UMI-tools or homotrimer correction.

While sequencing simulations can offer valuable insights, their real-world applicability may be limited by biases. To validate our homotrimer UMI error correction approach, we conducted experiments using a Common Molecular Identifier (CMI) attached to every captured RNA molecule (Supplementary Fig. 8). Having the same molecule attached to every RNA guarantees that, in the absence of errors, each transcript is only counted once. However, if errors are introduced into the CMI, transcripts will be overcounted. This provides a means for assessing the accuracy of library preparation and sequencing, as well as the impact of errors on the transcript counts (Supplementary Fig. 8b).

We attached the CMI to equimolar concentrations of mouse and human cDNA at the 3’ end, PCR amplified and split the sample for sequencing on Illumina, PacBio or ONT platforms. We calculated the hamming distance between the observed and expected CMI sequence to measure sequencing accuracy. Our results show that 73.36%, 68.08%, and 89.95% of CMIs were accurately called using Illumina, PacBio, and the latest kit14 ONT chemistry, respectively (Fig. 1h, Supplementary Fig. 9 and supplementary Fig. 10). Older ONT chemistry gave substantially lower accuracy (Supplementary Fig. 11), but the use of super accuracy basecalling led to substantial improvements (Supplementary Fig. 12). Using our homotrimeric error correction approach, we were able to correct 98.45%, 99.64% and 99.03% of CMIs for Illumina, PacBio and the latest ONT chemistry, respectively (Fig. 1h). We hypothesised that the lower accuracy of Illumina and PacBio when compared to ONT sequencing may be due to higher number of PCR amplification cycles during sequencing (e.g. bridge amplification and rolling circle amplification with Illumina and PacBio, respectively). In order to untangle the effect of sequencing and PCR errors, we subjected a CMI-tagged cDNA library to increasing cycles of PCR and then sequenced using ONT’s Minion platform. To minimise batch effects across our data and simultaneously evaluate sequencing accuracy independent of PCR amplification effects, we attached trimer sample barcodes during PCR amplification. We show a high degree of barcode accuracy and observed that homotrimer correction had minimal effect in improving this accuracy (Fig. 1i). Based on these results, it can be inferred that sequencing errors make a negligible contribution to the overall error rate. However, we observed a substantial increase in the number of errors within our CMIs with increasing PCR cycles (Fig. 1j). Our homotrimer approach was able to correct the majority of errors observed within the CMIs (Fig. 1k). This suggests that PCR is a significant source of UMI error. We next benchmarked homotrimer error correction against UMI-tools and found substantial improvements in error correction (Fig. 1l). Furthermore, using pairs of UMIs we were able to remove most human-mouse chimeric artefacts from our library. This dual UMI tagging approach was not only experimentally validated, but also confirmed our previous simulations (Supplementary Fig. 7), ultimately leading to improved accuracy and reliability of chimeric artefact removal (Fig. 1m).

Having demonstrated the ability to accurately correct PCR errors using homotrimers, we conducted an experiment in which we treated Ewing’s RM82 sarcoma cells with an inhibitor for the splicing kinase CLK1. This was done to induce splicing perturbations and to observe an exaggerated differential transcript effect, after sequencing by either ONT’s PromethION or Illumina platforms (Fig. 1n-p, Supplementary Fig. 13 and Supplementary Fig. 14). When we compared monomer UMI correction (i.e. the selection of a random monomer in the homotrimer and applying UMI-tools deduplication method) to our homotrimer correction methodology, we found differences in the number of differentially expressed genes and transcripts between splicing inhibition and control conditions. Specifically, for genes and transcripts, the difference was 7.8% and 11%, respectively (Fig. 1n-o). We also observed 4.7% discordant differentially expressed genes following Illumina sequencing (Supplementary Fig. 14c-d). We show that homotrimer error correction corrects PCR amplification errors that lead to increased UMI variability between samples on a per gene level (Fig. 1p). In addition, the homotrimer correction approach led to an increased fold enrichment of genes associated with gene ontology terms related to DNA replication and splicing (Supplementary Fig. 15), highlighting the improved accuracy of our method in identifying biologically relevant gene sets.

To understand how PCR errors contribute to single-cell sequencing errors we encapsulated JJN3 human and 5TGM1 mouse cells using the 10X Chromium system and performed reverse transcription followed by 10 PCR cycles. Subsequently, we divided the PCR product into two portions and performed additional PCR amplification, resulting in a combined number of PCR cycles of 20 or 25. We then prepared and sequenced these libraries using ONT’s PromethION platform, which does not perform PCR amplification as part of the sequencing process. After assigning cell barcodes (Fig. 2a) and filtering, clustering and annotating the cells (Fig. 2b), we observed unexpectedly that the library subjected to 25 cycles of PCR had significantly greater number of UMIs compared to the library that underwent 20 PCR cycles (Fig. 2c and Supplementary Fig. 16). This suggests that PCR errors may contribute to inaccurate counting of transcripts and an inflated UMI count. We next performed differential gene expression between 20 and 25 PCR cycles in both the mouse and human cells and identified 50 differentially expressed transcripts (Supplementary Table 2 and Supplementary Table 3). For example, transcripts ENSMUST00000034966 (Fig. 2d; Rpl4, ribosomal protein L4) and ENST00000532223 (Fig. 2e; IGLL5, immunoglobulin lambda), were identified as highly significant in this differential expression analysis, highlighting the contribution of PCR errors to inaccurate transcript counting.

**Figure 2:**
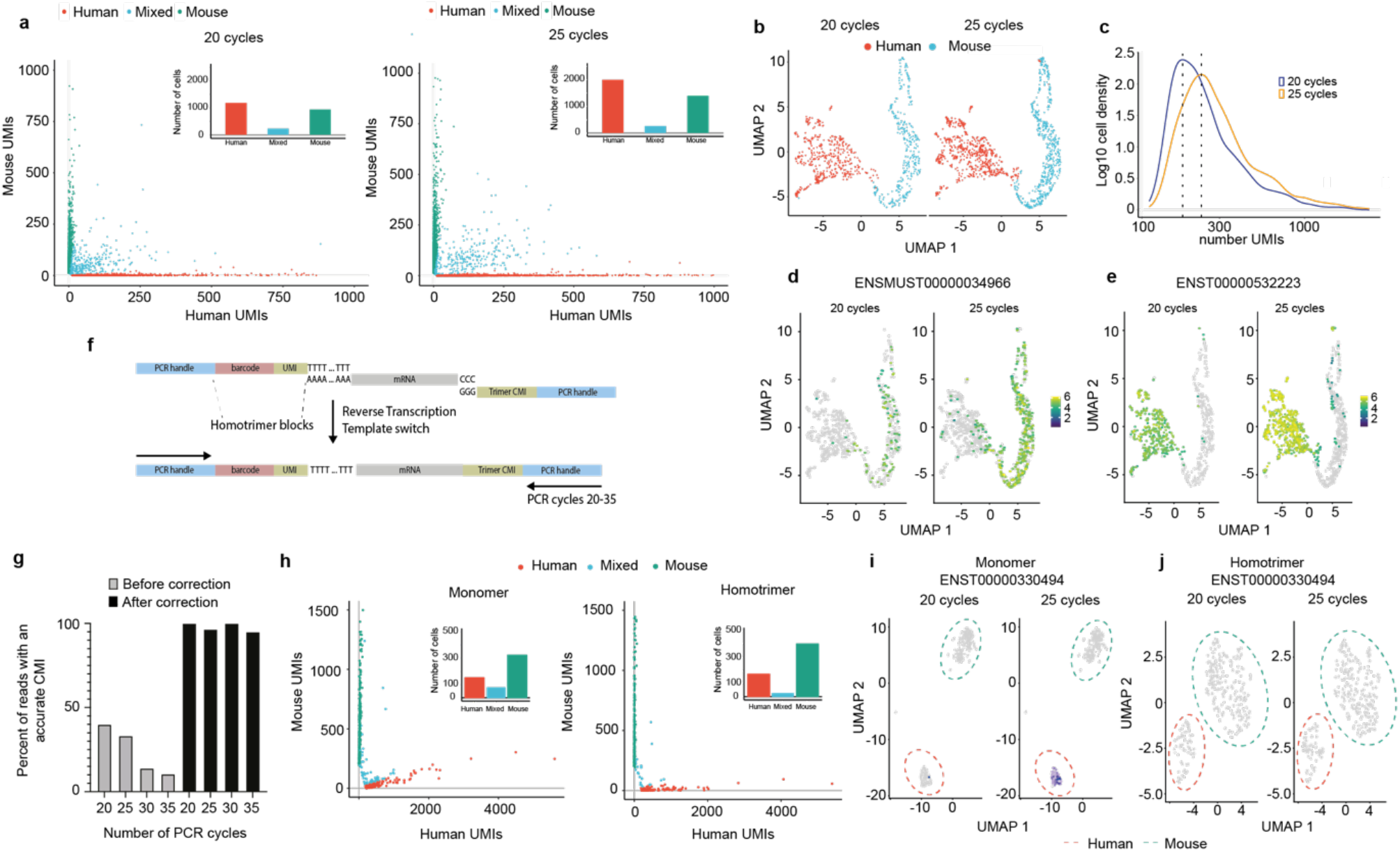
**a**, Human Jurkat and mouse 5TGM1 cells were mixed at a 50:50 ratio and approximately 3000 cells were taken for encapsulation and cDNA synthesis using 10X chromium followed by nanopore sequencing. **b**, A UMAP of 10X chromium data showing the integration, clustering and annotation of human and mouse cells following 20 and 25 cycles of PCR. **c**, A density plot for the 10X chromium data showing the log10 density of the number of UMIs following 20 and 25 cycles of PCR. The dotted line shows the maximum density for each condition. **d**, A UMAP showing the expression of ENSMUST00000034966 between libraries amplified following 20 and 25 PCR cycles. **e**, A UMAP showing the expression of ENST00000532223 between libraries amplified following 20 and 25 cycles of PCR. **f**, Human JJN3 and mouse 5TGM1 cells were mixed at a 20:80 ratio and approximately 500 cells were PCR amplified. A Schematic showing the homotrimer UMI drop-seq library preparation approach and template switching attachment of a homotrimer CMI to single-cell captured mRNAs. **g**, Drop-seq libraries were sequenced using the Flongle™ sequencing device, graphs show the percent of reads that have an accurate CMI following amplification of the same library using 20, 25, 30 and 35 cycles of PCR before and after homotrimer correction. **h**, barnyard plots showing the expression of mouse and human cells following 20 and 25 cycles of PCR and sequencing using a PromethION™ sequencing device. UMAP plot showing the transcript expression of ENST00000330494 following monomer based UMI-tools demultiplexing (**i**) and homotrimer based demultiplexing (**j**).

Next, we encapsulated JJN3 human and 5TGM1 mouse cells and performed reverse transcription and template switching that included a CMI, followed by 10 PCR cycles. Subsequently, we divided the PCR product into four portions and performed additional PCR amplification so that each library was subjected to 20, 25, 30 and 35 PCR cycles. The libraries were then sequenced using the ONT Minion platform. Our results indicate a decrease in the percentage of reads with accurate CMIs as the number of PCR cycles increases. Importantly, we show that homotrimer correction leads to 96-100% correction of CMI sequences (Fig. 2g and Supplementary Fig 17). This underscores the effectiveness of this approach in removing errors introduced by PCR. Subsequently, we sequenced the libraries that underwent 20 or 25 PCR cycles using ONTs PromethION platform (Supplementary Fig. 18). Our results show that by incorporating homotrimers within the barcode region an increase, albeit low (∼15%), in the numbers of cells recovered was achieved (Fig. 2h). Monomeric UMIs resulted in over 300 differentially regulated transcripts between the 20 and 25 cycle libraries (Fig. 2i and Supplementary Table 4). On the contrary, homotrimer correction found no significant differentially regulated transcripts (Fig. 2j), demonstrating the robustness of homotrimer UMIs to remove errors.

Our study underscores the crucial role of precise UMI counting in both bulk and single-cell sequencing, and proposes homotrimer UMIs as a reliable experimental solution to enhance the accuracy of absolute read counting. PCR amplification errors are the main source of UMI inaccuracy, but homotrimer UMIs can improve error correction and achieve near perfect sequencing of molecule counts. This study has implication for researchers conducting comparative and longitudinal sequencing studies, where precise UMI counting is crucial, yet its importance is often overlooked.

## Methods

### Cell lines and reagents

5TGM1, Jurkat and RM82 cell lines were cultured in complete RPMI medium. All parental cell lines were tested twice per year for mycoplasma contamination and authenticated by STR during this project. For cell culture experiments, SGC-CLK-1 (Structural Genomics Consortium) inhibitor was incubated with cells for 24 hours. DMSO was used as a negative control.

### Oligonucleotide synthesis

Homotrimer phosphoramidites were purchased as a custom product from Metkinen Chemistry (Finland) and reverse homotrimer phosphoramidites were a custom synthesis product from Chemgenes (USA). Solid-phase phosphoramidite oligonucleotide synthesis on Toyopearl HW-65S resin (Tosoh Biosciences, 0019815) was performed by ATDBio as described previously^12^, in the 5′–3′ direction (using reverse amidites), using a method adapted from Macosko et al ^16^. The sequence of the capture oligonucleotide is as follows: Bead-5’-[spacer]-TTTTTTTAAGCAGTGGTATCAACGCAGAGTACJJJJJJJJJJJJNNNNNNNNTTTTTTTTTTTTTTTTTTTTTT TTTTTTTT-3′, where ‘J’ indicates a nucleotide trimer block added via split and pool synthesis using reverse monomer phosphoramidites. ‘N’ indicates a degenerate trimer nucleotide (added using an equimolar mixture of the four reverse timer phosphoramidites). [spacer] is hexaethylene glycol, added using DMT-protected hexaethylene glycol phosphoramidite (HEG), and all the other bases are standard (monomeric) DNA bases, added using reverse phosphoramidites. AAGCAGTGGTATCAACGCAGAGTAC is the PCR handle.

Before oligonucleotide synthesis, capping was performed to reduce the initial loading of hydroxyl groups on the beads, by suspending the resin in a 1:1 mixture of Cap A (tetrahydrofuran:lutidine:acetic anhydride 8:1:1) and Cap B (tetrahydrofuran:pyridine:1-methylimidazole 8:1:1) at room temperature for 30 min. Oligonucleotide synthesis was then performed using an ABI 394 DNA synthesizer, using a modified 1 μmol synthesis cycle (with an extended coupling time of 5 min for monomer bases and 10 min for trimer bases. The capping step was omitted for the trimer bases in the UMI region and the poly-T region). The barcode was generated using 12 split-and-pool synthesis cycles. Before the first split-and-pool synthesis cycle, beads were removed from the synthesis column, pooled and mixed, and divided into four equal aliquots. The bead aliquots were then transferred to separate synthesis columns before three consecutive couplings with monomers reverse amidites. This process was repeated 11 times. Following the final split and pool cycle, the beads were pooled, mixed and divided between four columns, ready for the next part of the synthesis. An equimolar mixture of the four trimer phosphoramidites was used in the synthesis of the degenerate UMI (poly(N)) region, and (monomeric) T reverse amidite was used for the poly(T) tail. After oligonucleotide synthesis, the resin was washed with acetonitrile and dried with argon before deprotection in aqueous ammonia (r.t., 17h followed by 55 °C, 6 h). The beads were then washed with water followed by acetonitrile and dried with argon gas.

Template switch oligonucleotide was synthesized using standard phosphoramidites: 5’ - AAGCAGTGGTATCAACGCAGAGTNNNNNNNNNNGAATrGrGrG-3’. The oligonucleotides were PAGE purified and shipped lyophilized. Primers containing Common Molecular Identifiers (CMI) were synthesised by Sigma Aldrich (Burlington, USA) using the following sequences polyA oligonucleotide: 5’ - AAGCAGTGGTATCAACGCAGAGTACNNNNNNNNNNTTTTTTTTTTTTTTTTTTTTTTTTTTTTTT-3

### Generating bulk homotrimer UMI tagged cDNA

Total mRNA was isolated using a Quick-RNA MiniPrep kit (Zymo), following the manufacturers protocol. The RNA sample quality and quantity was measured using an RNA screen tape on the TapeStation (Agilent). cDNA synthesis was performed with modification to the SMART approach^17^. An oligo(dT) containing adaptor containing a homotrimer 30-base DNA sequence and a SMART primer sequence was used to initiate a reverse transcriptase reaction. Briefly, RNA was denatured at 72°C for 2 minutes and then reverse transcribed with Maxima H minus reverse transcriptase (2000 U) in a total volume of 50uL with the buffer, 1mM dNTPs, 2mM dithiothreitol (DTT) and 4% Ficoll PM-400. The reaction was performed for 90 minutes at 42 °C and then the enzyme was heat inactivated at 80 oC for 5 minutes. The library was then purified using 0.8X SPRI bead (Beckman Coulter) clean-up followed by PCR using the SMART PCR primer (AAGCAGTGGTATCAACGCAGAGT) before being purified using SPRI beads. To achieve a high concentration of cDNA the input was subjected up to 30 cycles of PCR amplification followed by a second cleanup. Optionally, 10ng of PCR product was subjected to 12 further cycles of PCR using primers that contained trimer sample barcodes (Supplementary Table 1). Finally, cDNA was quantified using a TapeStation (Agilent Technologies) using a DNA high-sensitivity D5000 tape before being split for Illumina or Oxford Nanopore library generation. To reduce PCR artefacts and improve sequencing return, we performed PCR using the primer 5—PCBio-TACACGACGCTCTTCCGATCT further 3-5 cycles of PCR

### ONT bulk RNA seq library preparation and sequencing

A total of 1,200ng of purified cDNA was used as a template for ONT library preparation. We used SQK-LSK-109, SQK-LSK112 and SQK-LSK114 (also referred to as ONT latest kit14 chemistry). Ligation sequencing kit, following the manufacturers protocol. Samples were sequenced using a minION™ device using R9.4.1 (FLO-MIN106D) or R10.4 (FLO-MIN112) flow cells. Barcoding using the Native Barcoding Amplicon kit (EXP-NDB104) was performed for RM82 cells treated with DMSO or CLK1 inhibitor treatment. These samples were sequenced using the PromethION™ sequencing platform on R9.4.1 FLO-PRO002 flow cells at the Deep Seq facility at the University of Nottingham.

### PacBio bulk RNA seq library preparation and sequencing

A total of 1,200ng of purified cDNA was used as a template for PacBio library preparation and sequencing at the Centre for Genomic Research at the University of Liverpool (https://www.liverpool.ac.uk/genomic-research/technologies/next-generation- sequencing/). cDNA was end repair and A tailed with T4 polynucleotide kinase (New England Biolabs). The sequencing library was prepared using SMRTbell Express Template Prep Kit 2.0 following the standard protocol. Sequencing was then performed on a sequel II using a Sequel II SMRT Cell 8M ion CCS mode, following the standard protocol. CCS reads were generated using CCS v6.3.0 (https://github.com/PacificBiosciences/ccs) using default settings.

### Illumina bulk RNA seq library preparation and sequencing

Purified cDNA was used as an input for the Nextera XT DNA library preparation kit (New England Biolabs). Library quality and size was determined using a TapeStation (Agilent Technologies) High Sensitivity D1000 tape and then sequenced on a NextSeq 500 sequencer (Illumina) using a 75-cycle High Output kit using a custom read1 primer (GCCTGTCCGCGGAAGCAGTGGTATCAACGCAGAGTAC). Read1 length was 30bp and read2 length was 52bp long.

### ONT bulk RNA sequencing analysis workflow

The data was processed using a custom pipeline *‘pipeline_count’* written using cgatcore and included within the TallyNNN repository^18^. Briefly, the quality of each fastq file was evaluated using fastqc toolkit ^19^ and summary statistics were collated using Multiqc^20^. We then identify the polyA associated UMI sequence by searching for the polyA region and reverse complementing the read if it does not appear in the correct orientation. The 30bp UMI is then identified upstream of the SMART primer by pattern matching for GTACTCTGCGTTGATACCACTGCTT. We then corrected for errors or remove the read based the number of UMI errors and then the UMI is added to the read name. Next, the TSO associated UMI is identified using the SMART primer sequence AAGCAGTGGTATCAACGCAGAGTAAT. The 30bp UMI sequence is then corrected for errors or remove the read based the number of UMI errors and then the UMI is added to the read name. Both the TSO and polyA associated UMIs and primer sequences are removed from the read sequence. For transcrip0t level analysis, the fastq file is then mapped against the transcriptome using minimap2 (v2.22) with the following settings: -ax map-ont -p 0.9 --end-bonus 10 -N 3. The resulting sam file was then sorted and indexed using samtools^21^. A custom script was then used to add the transcript name to the XT tag of the samfile for downstream counting by homotrimer deduplication, UMI-tools or mclUMI. For gene level analysis, the fastq data was mapped using minimap2 using the following setting: -ax splice -k 14 --sam-hit-only --secondary=no --junc-bed. The resulting sam file was then sorted and indexed followed by feature annotation using featurecounts (v2.0.1)^22^ using the following settings to generate an annotated bam file: featureCounts -a (gtf) -o (output) -R BAM. This bam file was then used for downstream counting by UMI-tools or mclUMI. The reference transcriptome and genomes used for the analysis were hg38_ensembl98 and mm10_ensembl88.

### Illumina bulk RNA sequencing analysis

The data was processed using a custom cgatcore written pipeline ‘pipeline_illumina’. Briefly, the UMIs contained in read1 were corrected based on homotrimer complementarity or were removed from the analysis depending upon a set error threshold. The paired fastq files were then mapped using hisat2 (v2.2.1)^23^ before features being counted using featureCounts using the following commands: featureCounts -a (gtf) -o (output) -R BAM. The resulting XT tagged bam file was then used for downstream counting using homotrimer deduplication, UMI-tools or mclUMI. Basic homotrimer deduplication, where a “majority vote” of each homotrigram is performed, can also be ran prior to UMI-tools and mclUMI by modifying the configuration file according to the documentation.

### UMI-tools deduplication

Following gene or transcript level mapping, the UMI was extracted from the read. Since UMI-tools was not designed to correct homotrimer sequences, we collapsed the UMI into a single nucleotide sequence by selecting the first base within each of the individual trimers. Reads were then deduplicated using the directional method using the command: umi_tools count – per-gene –gene-tag=XT.

### Homotrimer deduplication

Following gene or transcript level mapping, the UMI was extracted from the read and collapsed into single nucleotide sequence using the majority vote approach where applicable or resolve inconsistencies through a combinatorial optimization scheme otherwise. Briefly, reads were first filtered to exclude reads in which there were more than 3 errors in the UMI sequence. For UMI sequences where each trimer contains at least two identical nucleotides, a majority vote was then performed to collapse the trimer into a monomer. If at least one trimer is inconclusive and contains three different nucleotides, we no longer treat each UMI sequence independently when collapsing trimers into monomers. Instead, we select one of the nucleotides in each trimer block to achieve maximal consistency between duplicates, i.e. to minimize the number of distinct collapsed UMI sequences. We formulate this task as a set cover problem for each gene as follows^24^. Let *S* be the set of sequenced homotrimer UMIs of a given gene (in a given cell). For *s* ∈ *S* let *C*(*s*) denote the set of collapsed UMIs that can be obtained by combining single nucleotides that occur in each trimer block of *s*. Each such collapsed sequence *c* ∈ *C*(*s*), for some *s* ∈ *S*, can explain potentially multiple homotrimer UMIs *s*^′^ if *c* is also contained in *C*(*s*^′^). We therefore include one subset *S*_*c*_ ⊆ *S* for each *c* ∈ ⋃_{*s*∈*S*}_ *C*(*s*) that contains all *s* ∈ *S* for which *c* ∈ *C*(*s*). The collection of sets *S*_*c*_ of smallest cardinality that together include (“cover”) all sequenced UMIs in *S* therefore corresponds to the smallest set of collapsed UMIs that explain all *s* ∈ *S*. To find this smallest set of collapsed UMIs we employ a greedy algorithm that starts from the empty set and in each iteration adds the subset *S*_*c*_ (i.e. collapsed UMI *c*) that explains the largest number of yet unexplained sequenced UMIs. The solution returned by this algorithm is guaranteed to be within a logarithmic factor of the optimal solution^24^. In our experiments, the solution of the greedy approach was identical to the optimal solution for more than 90% of the genes. We computed the optimal solution using an integer linear programming approach, where decision variables model the inclusion or exclusion of sets *S*_*c*_ and linear inequalities enforce each sequenced UMI to be covered by at least one such set, i.e. to be explained by at least one collapsed UMI.

### Settings for simulated UMIs

We simulated UMI data of length 30 (10 blocks of nucleotide trimers) to confirm the accuracy of our UMI correction methodology by using the ResimPy tool. We mimicked the PCR amplification and sequencing errors seen with ONT sequencing, as this sequencing methodology suffers from indels and basecalling errors more frequently than PacBio or Illumina sequencing. UMIs were generated following an approach that was first described by UMI-tools^10^. Briefly, we simulated homotrimer blocks of UMIs at random, with an amplification rate (-ampl_rate) ranging between 0.8-1.0 and then simulated PCR cycles so that each UMI was duplicated to the probability of amplification. PCR errors were then randomly added and assigned new probabilities of amplification. A defined number of UMIs were randomly sampled to simulate sequencing depth and sequencing errors introduced with a specified probability. Finally, errors were detected by assessing the complementarity of homotrimers across the full UMI sequence. If no errors are detected, then the homotrimers are collapsed into single nucleotide bases. However, if errors are identified then collapsing into single nucleotides is performed using the most common nucleotide within the trimer. If a most common nucleotide cannot be determined, then a single nucleotide is selected at random for collapsing. The following values were used as values within our simulations. Sequencing depth 10-400; number of UMIs 10-100 (-umi_num); UMI-length 6 – 16 (-umi_len); PCR error rate 1×10^−3^ – 1×10^−5^ (-seq_err); sequencing error rate 1×10^−1^ – 1×10^−7^ and number of PCR cycles 4-12 (-pcr_num); permutation tests 10-100 (-perm_num).

### ResimPy – Simulating chimeric artefacts in UMI sequences

We developed a UMI simulation package called ResimPy. The number of UMI sequences *m* to be amplified at PCR *i* in ResimPy is described as the Galton-Watson branching (GWB) process

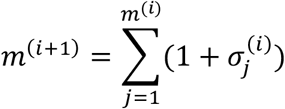

where 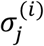 is the Kronecker symbol to represent the presence or absence of the *jth* sequence at PCR *i* with an amplification rate *α*, given by

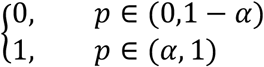

We used a binomial distribution *Binom*(*m*^(*i*)^, *α*) to generate *m*^(*i*+1)^. The 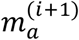 UMI sequences were chosen from its preceding PCR cycle *i* based on a uniform distribution *U*(0, *m*^(*i*)^). To simulate PCR errors, we also implemented another GWB process. The total number of base errors 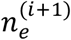 at PCR cycle *i* + 1 is modelled by

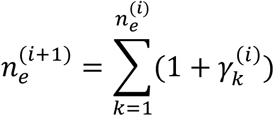

where 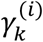 is the Kronecker symbol to indicate a binary state (*i*.*e*., erroneous or correct) of a position of sequences at PCR cycle *i*,

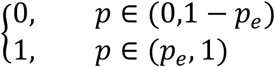

where *p*_*e*_ is the PCR error rate. We used a negative binomial distribution *NBinom*(*q* × *n*^(*i*)^, *q*) to generate 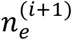, where *q* = 1 − *p*_*e*_. These 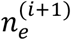 sequence positions were then randomly sampled based on a uniform distribution *U*(0, *n*^(*i*+1)^) where *n*^(*i*+1)^ represents the total number of bases at PCR cycle *i* + 1. Each PCR base error determined above is finally substituted by one of the remaining three types of bases to be drawn from a uniform distribution *U*(0,3). Likewise, we adopted the same way to simulate sequencing errors.

We simulated reads mixed with chimeric artefacts using the *resimpy_umi_transloc* module in Resimpy. Briefly, a total of 50 reads were generated, each attached with a UMI included at the 3’ end and a UMI at the 5’ end. Each randomly simulated UMI of homotrimer blocks was 30bp in length. We simulated UMIs with at least a 3-base edit distance away from one another. The chimeric frequency was set to 2% of the total read count. The number of wrongly synthesized bases due to DNA polymerase errors and sequencing errors were estimated from our bulk data using negative binomial models as given in the section above. All other parameters for simulation were kept the same as the simulation of UMIs as above.

### Removal of 5’ and 3’ UMI tagged chimeric artefacts

To remove chimeric artefacts we developed umiRarity to detect chimeric artefacts through combining UMIs at 3’ and 5’ ends into one merged sequence. We leverage the fact that chimeric artefacts will generate UMI pairs at a significantly lower frequency than real translocations. The combination of UMIs at both ends can further bring the frequency down to a level which is lower than that only using UMI at either 5’ or 3’ end. To this end, we counted the combined homotrimer UMIs and also the combined monomer UMIs corrected from their homotrimer UMIs. We then set a count threshold to remove reads tagged by the combined UMIs below this threshold and retain reads above this threshold. The removed reads are considered as chimeric artefacts and the remaining reads are classified as true translocated reads. Methodological details of the chimeric artefact detection process are illustrated in Supplementary Fig. 5. The performance of chimeric artefact removal was evaluated by comparisons between the percentage of successfully removed chimeric artefacts and the percentage of sacrificed true reads, and between the percentage of successfully removed chimeric artefacts and the percentage of successfully detected PCR duplicates. The count thresholds used in our study range from 1-5. The umiRarity method is available as an independent module in our package mclUMI, which can be accessed by ‘mclumi dechimeric -m dc_by_cnt’.

### Common Molecular Identifiers (CMI) and error evaluation in bulk sequencing

To measure the error rate and evaluate the accuracy of our UMIs following library preparation and sequencing we synthesised a common sequence (GGGAAACCCTTTGGGCCCTTTAAACCCTTT) in replacement of a UMI to our polyA capture oligonucleotide. Following sequencing the CMI sequence was identified upstream of the SMART primer by pattern matching for GTACTCTGCGTTGATACCACTGCTT. The accuracy of our CMI was then determined by comparing the expected synthesised sequence to the extracted CMI sequence. The percent of CMI that show full complementary with the expected sequence were counted and the number of errors were determined for the inaccurate CMIs.

### Comparison between UMI-tools and homotrimer CMI deduplication methods

After mapping the reads to the reference genome at the gene level, we processed the data using two different strategies: UMI-tools and homotrimer deduplication. For homotrimer deduplication, we used the full length of the CMI sequence, while for UMI-tools we collapsed the CMI into a monomer by selecting the first base for each trimer block. The inclusion of the CMI sequence to our reads provides an experimental ground truth with which to evaluate the accuracy of each deduplication strategy. To assess the accuracy of the final deduplicated counts, we compared them to the expected ground truth CMI gene count of 1.

### Identification of fusion transcripts within ONT sequencing data

Following gene level mapping of the ONT sequencing data, the sam file was filtered using samtools to remove all non-primary alignments and supplementary alignments. mclUMI was then ran to remove the chimeric artefacts from true genomic chimeric fusions using the following settings: mclumi dechimeric -m dc_by_cnt -ibam (infile) -tcthres 5 -obam dechimeric.bam -obam_c chimerical.bam. Chimeric reads were identified based upon the sam flag tag SA. All chimeric fusions were then annotated using the bed file of genes and genomic coordinates followed by filtering and then counting using pysam and the collections.Counter module in Python.

### 10X Chromium library preparation

We prepared a single-cell suspension using JJN3 and 5TGM1 cells using the standard 10X Genomics chromium protocol as per the manufacturer’s instructions. Briefly, cells were filtered into a single-cell suspension using a 40 μM Flomi cell strainer before being counted. We performed 10X Chromium library preparation following the manufacturers protocol. Briefly, we loaded 3,300 JJN3:5TGM1 cells at a 50:50 split into a single channel of the 10X Chromium instrument. Cells were barcoded and reverse transcribed into cDNA using the Chromium Single Cell 3’ library kit and get bead v3.1. We performed 10 cycles of PCR amplification before cleaning up the library using 0.6X SPRI Select beads. The library was split and a further 20 or 25 PCR cycles were performed using a biotin oligonucleotide (5-PCBio-CTACACGACGCTCTTCCGATCT) and then cDNA was enriched using Dynabeads™ MyOne™ streptavidin T1 magnetic beads (Invitrogen). The beads were washed in 2X binding buffer (10mM Tric-HCL ph7.5, 1mM EDTA and 2M NaCl) then samples were added to an equi-volume amount of 2X binding buffer and incubated at room temperature for 10 mins. Beads were placed in a magnetic rack and then washed with twice with 1X binding buffer. The beads were resuspended in H_2_O and incubated at room temperature and subjected to long-wave UV light (∼366 nm) for 10 minutes. Magnetic beads were removed, and library was quantified using the Qubit™ High sensitivity kit. Libraries were then prepared before sequencing.

### Dropseq library preparation

Single-cell capture and reverse transcription were performed as previously described^16^. Briefly, JJN3 and 5TGM1 cells (20:80 ratio) were filtered into a single-cell suspension using a 40 μM Flomi cell strainer before being counted. Cells were loaded into the DolomiteBio Nadia Innovate system at a concentration of 310 cells per μL. Custom synthesised beads were loaded into the microfluidic cartridge at a concentration of 620,000 beads per mL. Cell capture was then performed using the standard Nadia Innovate protocol according to manufacturer’s instructions. The droplet emulsion was then incubated for 10 mins before being disrupted with 1H,1H,2H,2H-perfluoro-1-octanol (Sigma) and beads were released into aqueous solution. After several washes, the beads were subjected to reverse transcription. Prior to PCR amplification, beads were treated with ExoI exonuclease for 45 min. PCR amplification was then performed using the SMART PCR primer (AAGCAGTGGTATCAACGCAGAGT) and cDNA was subsequently purified using AMPure beads (Beckman Coulter). The library was split and a further 20 or 25 PCR cycles were performed using a biotin oligonucleotide (5—PCBio-TACACGACGCTCTTCCGATCT) and then cDNA was enriched using Dynabeads™ MyOne™ streptavidin T1 magnetic beads (Invitrogen). The beads were washed in 2X binding buffer (10mM Tric-HCL ph7.5, 1mM EDTA and 2M NaCl) then samples were added to an equi-volume amount of 2X binding buffer and incubated at room temperature for 10 mins. Beads were placed in a magnetic rack and then washed with twice with 1X binding buffer. The beads were resuspended in H_2_O and incubated at room temperature and subjected to long-wave UV light (∼366 nm) for 10 minutes. Magnetic beads were removed, and library was quantified using the Qubit™ High sensitivity kit. Libraries were then prepared for sequencing.

### Bulk and single-cell library preparation and ONT sequencing

A total of 500 ng of single-cell PCR input was used as a template for ONT library preparation. Library preparation was performed using the SQK-LSK114 (kit V14) ligation sequencing kit, following the manufacturers protocol. Samples were then sequenced on either a Flongle™ device or a PromethION™ device using R10.4 (FLO-PRO114M) flow cells.

### 10X analysis workflow

To process the 10X chromium data, we wrote a custom cgatcore pipeline (https://github.com/cribbslab/TallyNNN/blob/main/tallynnn/pipeline_10x.py)^18^. We first determined the orientation of the reads and if a polyT sequence was detected we reverse complemented the read. Next, we identified the barcode and umi based on the pairwise alignment of the sequence AGATCGGAAGAGCGT and AAAAAAAAA and identified the sequence between these alignments. We next removed reads that were greater or equal to 28 bp and isolated the barcode as the first 16 bp and the UMI the following 12 bp. The barcode and UMI sequence were then appended to the name of the fastq read using the underscore delimiter. Next, to remove barcode errors we parsed the barcodes from each read in the fastq file and then selected the most common barcode sequences using the number of expected cells in our library as the threshold. Next, for every read in the fastq file we then identified the closest barcode match for each read, allowing for two mismatches. Mapping was performed using minimap2 (v2.22)^25^, with the following settings: -ax splice -uf –MD –sam-hit-only –junc-bed and using the reference transcriptome for human hg38 and mouse mm10. The resulting bam file was sorted and indexed before adding the transcript name to the XT tag within the bam file. Counting was then performed using UMI-tools –method=unique before being converted to a market matrix format. Raw transcript expression matrices generated by UMI-tools count were processed using R/Bioconductor (v4.0.3) and the Seurat package (v3.1.4). Transcript matrices were cell-level scaled and log-transformed. The top 2000 highly variable genes were then selected based on variance stabilising transformation which was used for principal component analysis (PCA). Clustering was performed within Seurat using the Louvain algorithm. To visualise the single-cell data, we projected data onto a Uniform Manifold Approximation and Projection (UMAP).

### Drop-seq analysis workflow

To process the drop-seq data, we wrote a custom cgatcore pipeline (https://github.com/cribbslab/Bulk-TallyNNN)^18^. We followed the workflow previously described for identifying barcodes and UMIs using scCOLOR-seq sequencing analysis^12^. Briefly, to determine the orientation of our reads, we first searched for the presence of a polyA sequence or a polyT sequence. In cases were the polyT was identified, we reverse complemented the read. We next identified the barcode sequence by searching for the polyA region and flanking regions before and after the barcode. The trimer UMI was identified based upon the primer sequence GTACTCTGCGTT at the TSO distal end of the read, allowing for two mismatches. Barcodes and UMIs that had a length less than 48 base pairs were filtered. To conduct monomer-based analyses, a random base was selected from each homotrimer in the UMI or CMI and collapsed into a monomer. Homotrimer UMI correction was performed following mapping using minimap2 (v2.22)^25^. Mapping settings we as follows: -ax splice -uf – MD –sam-hit-only –junc-bed and using the reference transcriptome for human hg38 and mouse mm10. The resulting sam file was sorted and indexed using samtools^21^. For monomer UMI, counting was performed using UMI-tools before being converted to a market matrix format. For homotrimer UMI correction, the counting was performed using the script *greedy*.*py* within the TallyNNN repository. Raw transcript expression matrices generated by UMI-tools count and *greedy*.*py* were processed using R/Bioconductor (v4.0.3) and custom scripts were used to generate barnyard plots showing the proportion of mouse and human cells. Transcript matrices were cell-level scaled and centre log ratio transformed. The top 3000 highly variable genes were then selected based on variance stabilising transformation which was used for principal component analysis (PCA). Clustering was performed within Seurat using the Louvain algorithm. To visualise the single-cell data, we projected data onto a Uniform Manifold Approximation and Projection (UMAP).

## Data availability

Sequencing data has been deposited to GEO under the accession number GSE218899.

## Code availability

Source data is provided with this manuscript. All custom pipelines used within this analysis are available on github (https://github.com/cribbslab/TallyTriN). mclUMI is also available on github (https://github.com/cribbslab/mclumi). ResimPy is available on github (https://github.com/cribbslab/resimpy).

