## Supplementary Figures for "Correcting PCR amplification errors in unique molecular identifiers to generate absolute numbers of sequencing molecules"

Supplementary Figure legends

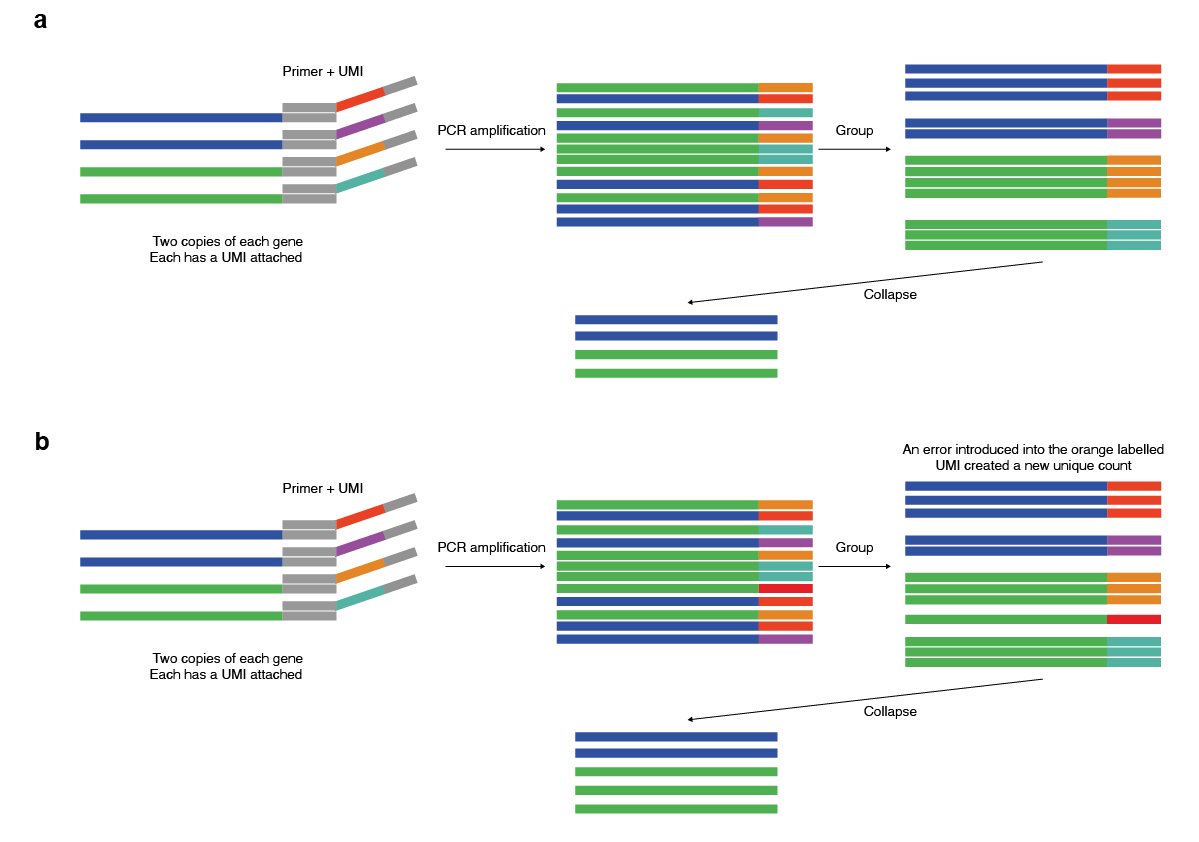

**Figure 1: UMI errors from sequencing or PCR increase UMI counting.**

**a**, Ideal UMI collapsing example: This figure illustrates an ideal scenario where transcripts (two blue and two green) are levelled with unique molecular identifier barcodes (UMIs) and PCR amplification is performed. Due to PCR amplification bias, longer transcripts have a lower amplification rate than shorter ones. The sequenced reads are then grouped together based on the set of UMIs and then collapsed within those groups to match the original number of transcripts. **b**, Increased UMI counting occurs due to errors: In real situations, errors occur during PCR amplification and sequencing, which can lead to increased UMI counts. As shown in this example, an error occurs within one of the UMIs which results in a higher count of unique UMIs than the actual number of input transcripts in the final library. This phenomenon can affect downstream analysis and should be taken into consideration otherwise they will lead to false positives when performing differential expression analysis.

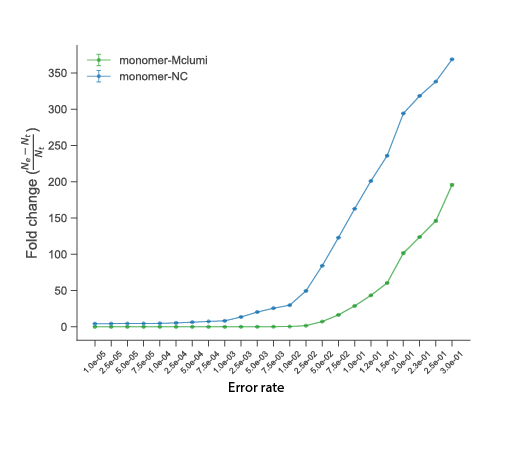

**Figure 2: Computational UMI demultiplexing strategies alone are insufficient to correct errors.**

To assess the efficiency of computational UMI demultiplexing strategies in correcting sequencing errors, we simulated monomer base UMIs with increasing error rates and calculated the fold change between the ground truth and the output of applying MCL-umi error correction. Our results show that even with the application of MCL-umi error correction, computational strategies alone are unable to fully correct sequencing errors, as evidenced by the increasing deviation from the ground truth with higher error rates.

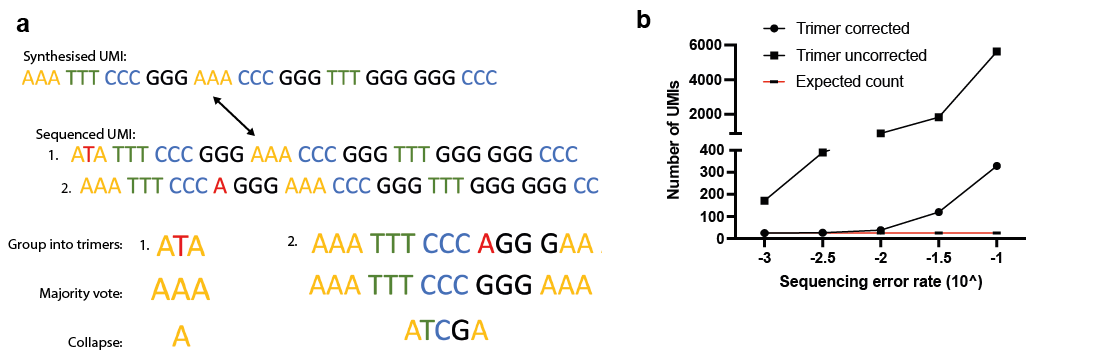

**Figure 3: Improved UMI deduplication using homotrimer blocks of nucleotides.**

**a**, Homotrimer blocks of nucleotides are used to synthesise UMIs, which enables efficient error correction through a majority vote between the trimer blocks. This approach does not require knowledge of the original oligonucleotide, and the trimers are resilient to base pair errors (example 1) and insertions/deletions (example 2). Errors are removed before collapsing the sequence to a single-base and then performing downstream analyses. **b**, We simulated UMIs with increasing error rates were modelled the correction of trimer sequences using the majority vote approach as described in **a**. To handle homotrimer errors more robustly, we subsequently developed a new model, which is described within the methods section “homotrimer correction”.

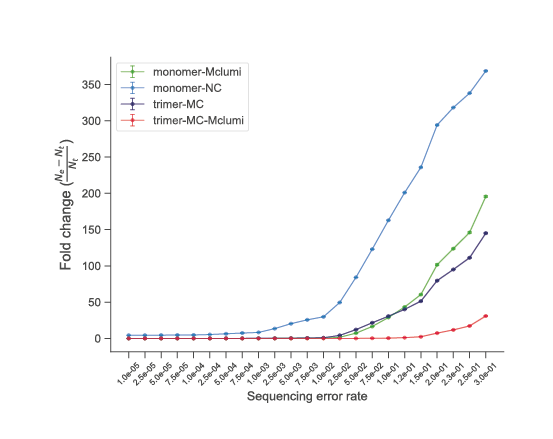

**Figure 4: Comparison of Homotrimer and monomer UMI for demultiplexing.**

We evaluated the effectiveness of homotrimer UMIs versus monomer UMIs for demultiplexing by simulating base UMIs with increasing error rates and calculated the fold change between the ground truth and the output of applying computational error correction. Our findings demonstrate that homotrimer UMIs outperform both uncorrected monomer UMIs and monomer UMIs corrected using computational methods. At very high error rates, the demultiplexing accuracy of homotrimer UMIs can be further improved by combining them with computational error correction techniques such as MCL-umi.

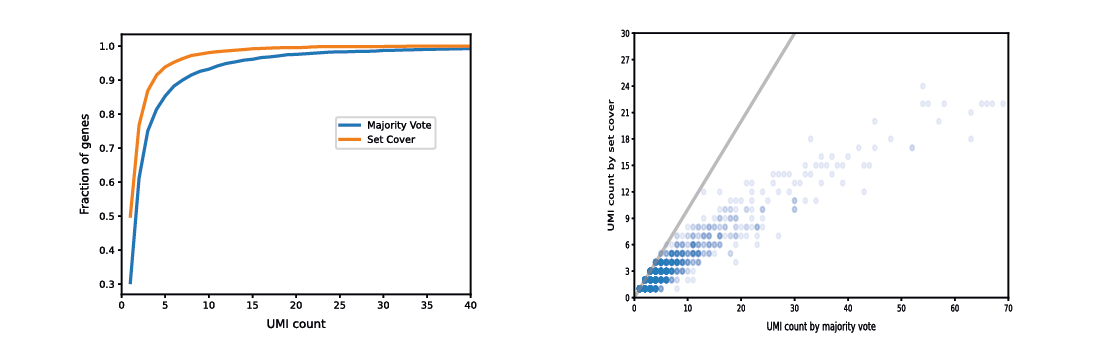
**Figure 5: The majority vote method is improved using a set coverage solution.**

**a**, A cumulative plot showing the fraction of genes that have been collapsed to a UMI count of less than or equal to the value shown on the x-axis by either the majority vote approach or the set cover-based optimization method. Only genes with at least 2 mapped reads were considered (n=3,428). Maximal UMI counts returned by majority vote and set cover optimization are 245 and 72, respectively. **b**, A scatter plot comparing UMI counts obtained using the majority vote approach (x-axis) to counts returned by the greedy set cover algorithm (y-axis). Only genes with at least 2 mapped reads were considered (n=3,428). To simplify visualization, we excluded genes with large majority counts (x,y) = (83, 22), (91, 24), (96, 31), (106, 32), (113, 33) and (245, 72).

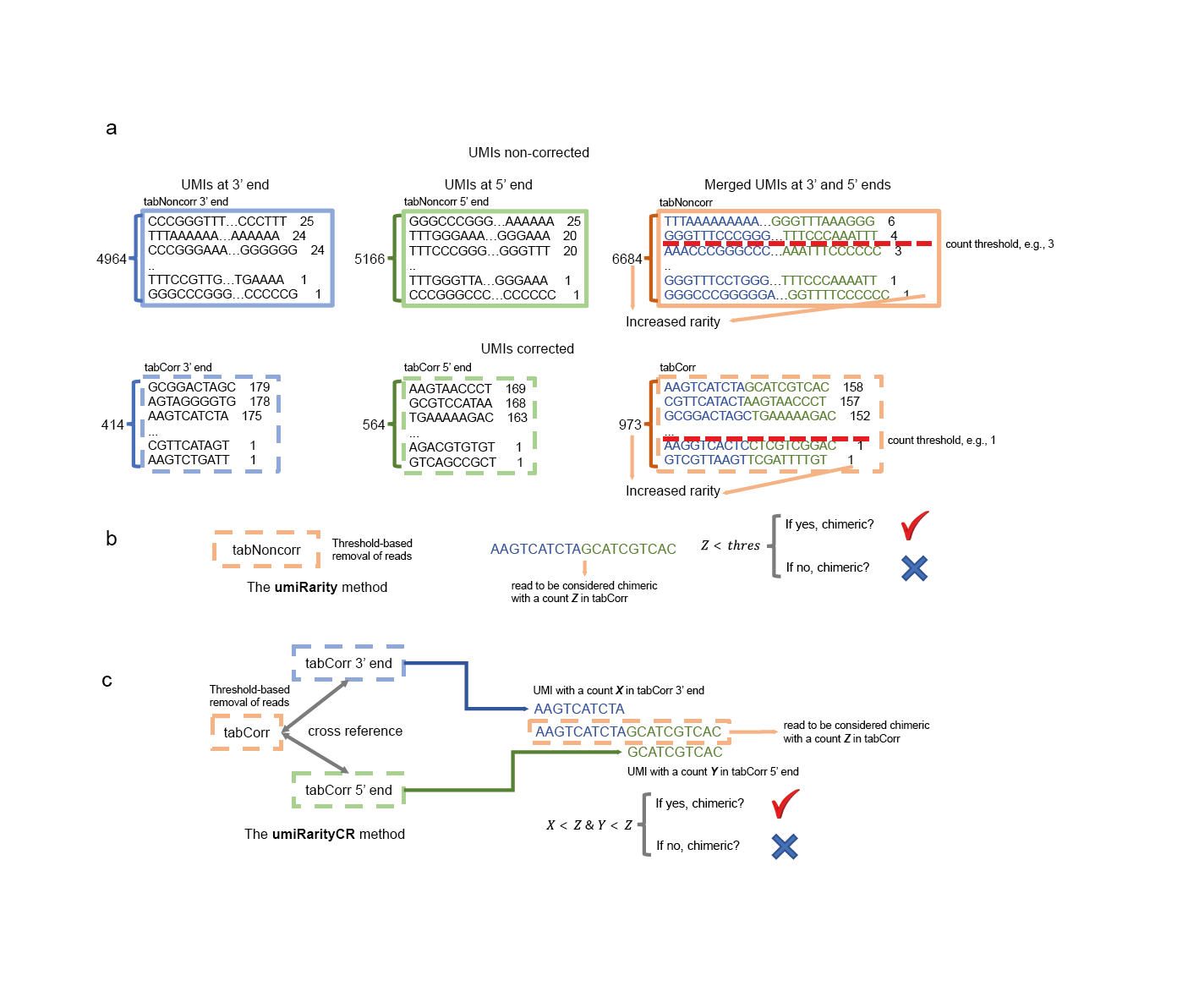

**Figure 6: Illustration of removal of chimeric reads using the Rarity and RarityCR methods.**

**a**, UMI count tables at a given genomic locus. It includes a count table of UMIs at 3’ end, a count table of UMIs at 5’ end, and a count table of UMIs obtained by merging UMIs at 3’ end 5’ ends. Non-corrected UMIs represent UMIs of trimer blocks in their original form, while corrected UMIs represent UMIs of collapsing trimer blocks. **b** and **c** show the Rarity and RarityCR methods, respectively. The Rarity and RarityCR methods are incorporated into the MCL-umi software tool.

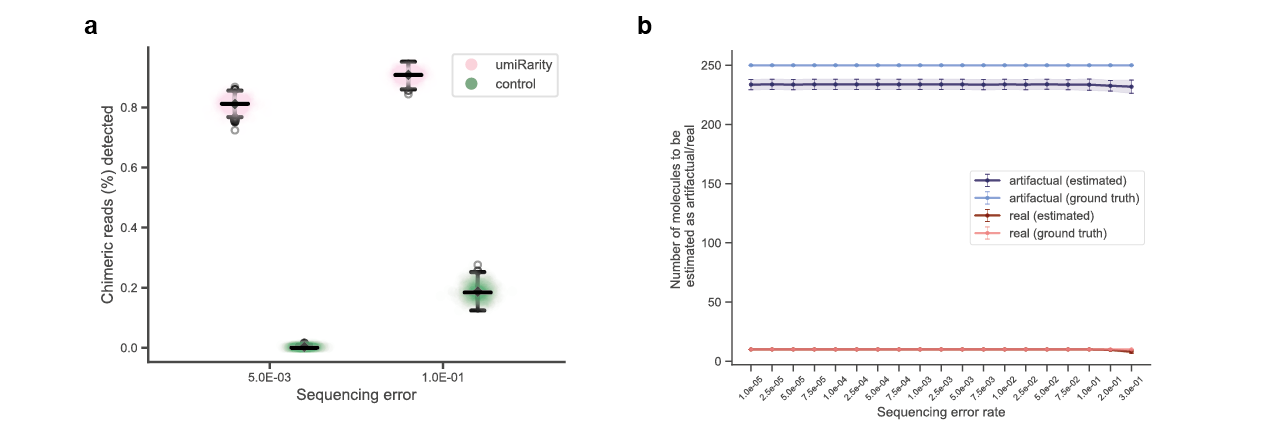

**Figure 7: Simulated data demonstrates the effectiveness of using dual UMIs for chimeric artefact removal.**

To evaluate the effectiveness of dual UMIs in removing chimeric artifacts, we simulated these events and tested the ability of dual UMIs attached to each cDNA end to accurately identify and remove these artifacts. During simulated PCR amplification, we assumed that there was a 2% probability of artefactual translocations occurring between two reads. Our goal was to evaluate the efficacy of using trimer UMIs at both ends to detect these artefactual translocations. We collapsed the trimer UMIs at both ends of each read into monomer UMIs and compared them to the original pairs of UMIs made during library preparation. A read was considered a real chromosomal translocation if its collapsed UMI pair was no more than 2 edit distance away from any UMI pairs in the prepared library; otherwise, it was considered a real PCR artifact. **a**, We evaluated the ability to detect chimeric artefacts using umiRarity and compared this to detecting artefacts only considering using UMIs at the 3’ end of the transcript (referred to as “control”). We show that considering UMIs at both ends is necessary for improving the performance of detecting chimeric artefacts.

**b**, We found that our threshold-based match strategy accurately identified the real and artefactual molecules, with the number of estimated molecules closely mirroring the ground truth. Our simulation suggests that the use of trimer UMIs can effectively remove PCR-amplified real and artefactual translocation molecules, which is an important consideration for translocation detection.

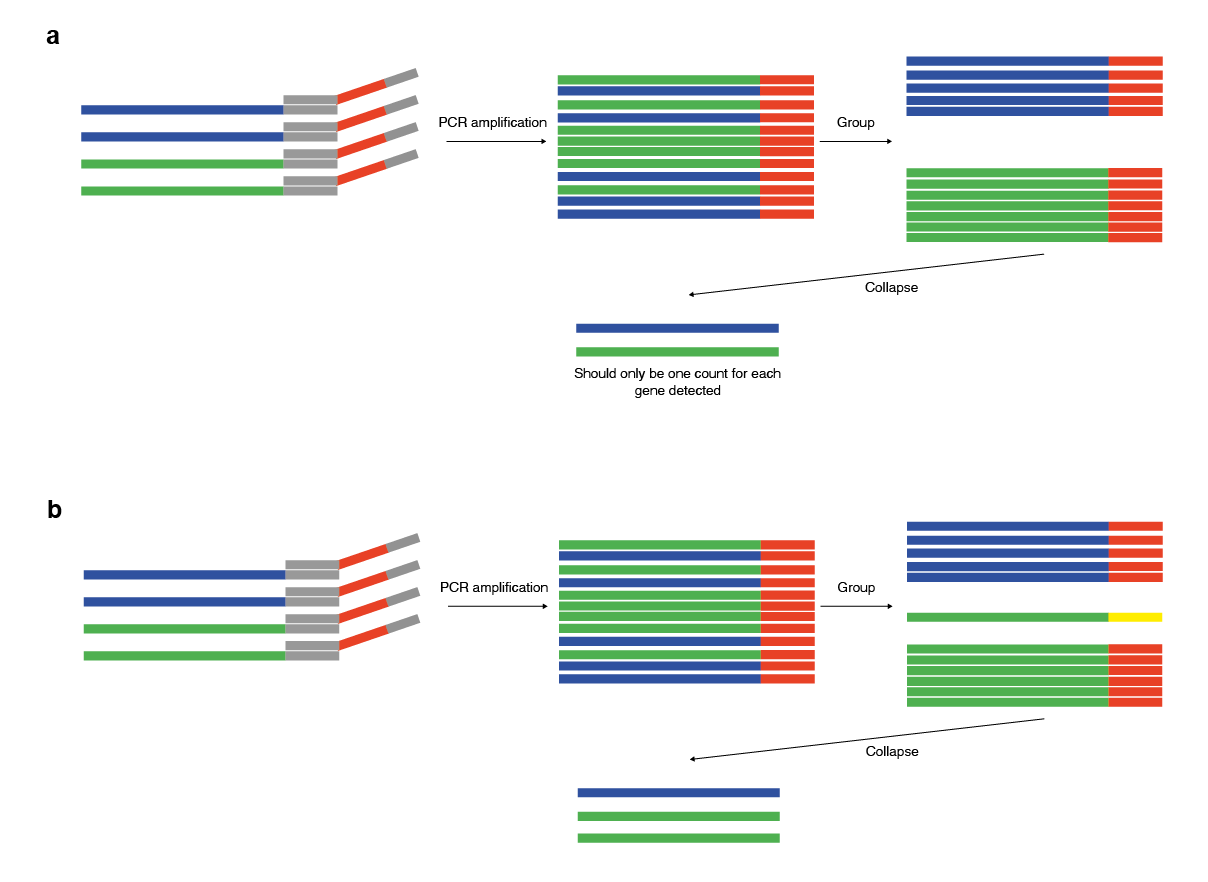

**Figure 8: Empirical evaluation of transcript counting with Common Molecular Identifiers (CMIs).**

**a**, An Ideal CMI collapsing example: In this scenario, transcripts (two blue and two green) are levelled with a common molecular identifier barcode (CMIs; labelled as red) and amplified via PCR. During transcripts grouping, all transcripts are labelled with the same common sequence. Therefore, following demultiplexing, each transcript should receive a count of one for every instance of detection. **b**, Increased counts result from the introduction of errors: This figure illustrates the effect of the errors within the CMI sequence. Any error introduced during PCR or sequencing creates a new CMI (labelled as yellow), resulting in an increase in transcript counts. This allows empirical evaluation of the effect of errors on the counting of transcripts, providing valuable insights into the accuracy of transcript quantification.

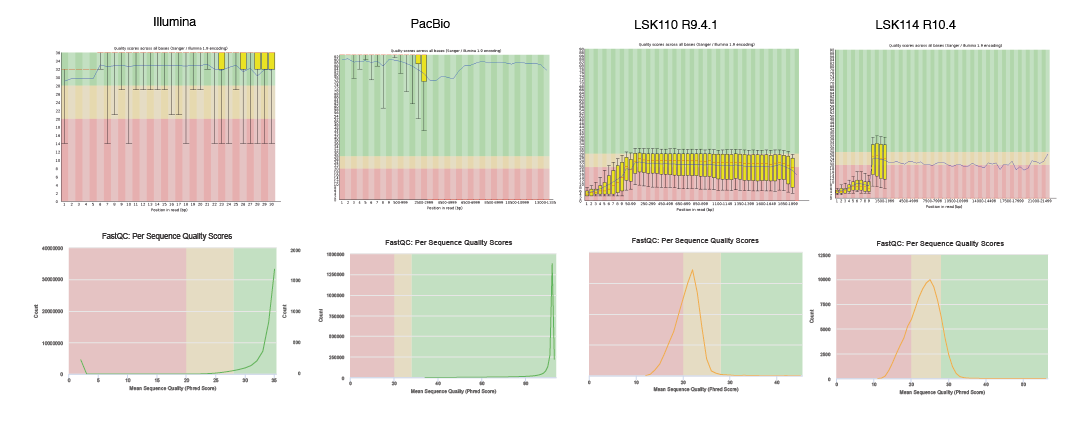

**Figure 9: Read quality control following sequencing by Illumina, PacBio and ONT.**

The read quality outputs from FASTQC for Illumina, PacBio and ONT (old chemistry: LSK110 R9.4.1 and new kit14 chemistry LSK114 R10.4).

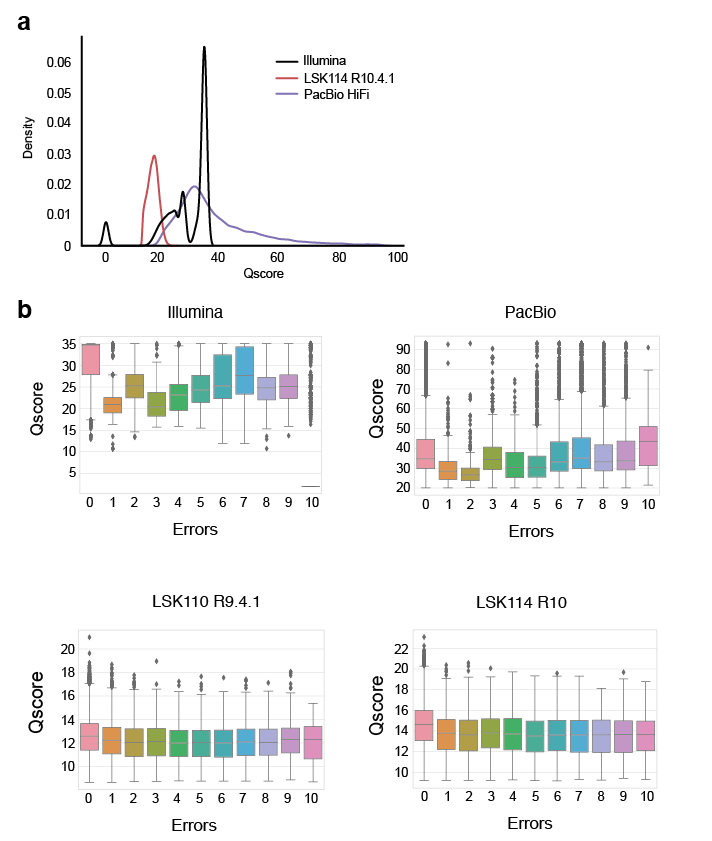

**Figure 10: Quality score following sequencing by Illumina, PacBio and ONT.**

The Qscore of each read as a relationship between the number of errors measured within the CMI, sequencing across Illumina, PacBio and ONT (old chemistry: LSK110 R9.4.1 and new kit14 chemistry LSK114 R10.4) technologies. **a**, Qscore represented as a density plot for the different sequencing platforms. **b**, The Qscore relationship with the number of errors detected within the CMI across the different sequencing platforms.

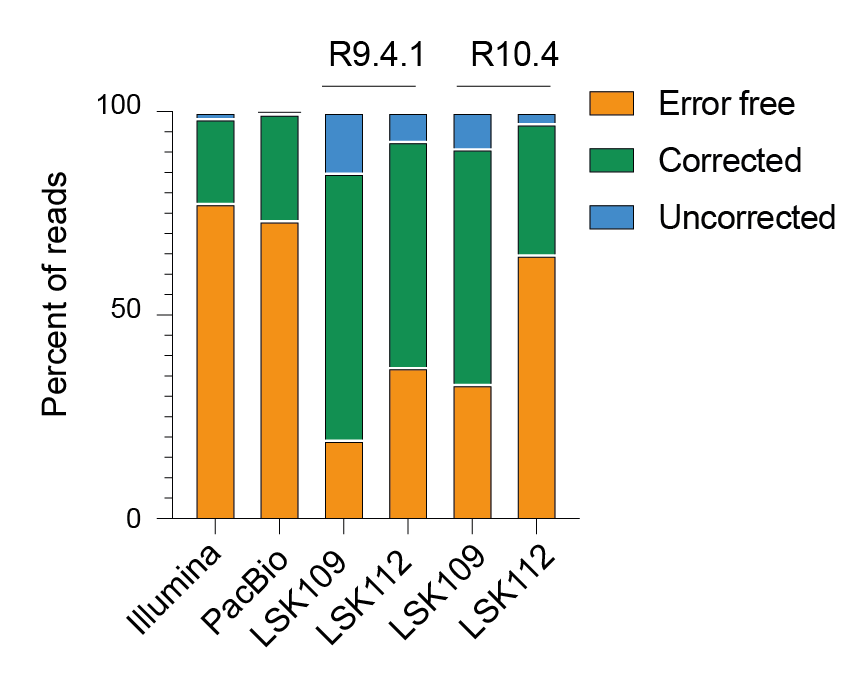

**Figure 11: Evaluation of CMI accuracy across Illumina, PacBio and legacy ONT chemistry.**

The accuracy of sequencing was evaluated for legacy ONT chemistry, we measured the percentage of CMIs with a Hamming distance between the expected and the sequenced CMI. The results are shown for Illumina, PacBio, and ONT legacy chemistry sequencing. Data from Illumina and PacBio is the same as in Fig. 1h.

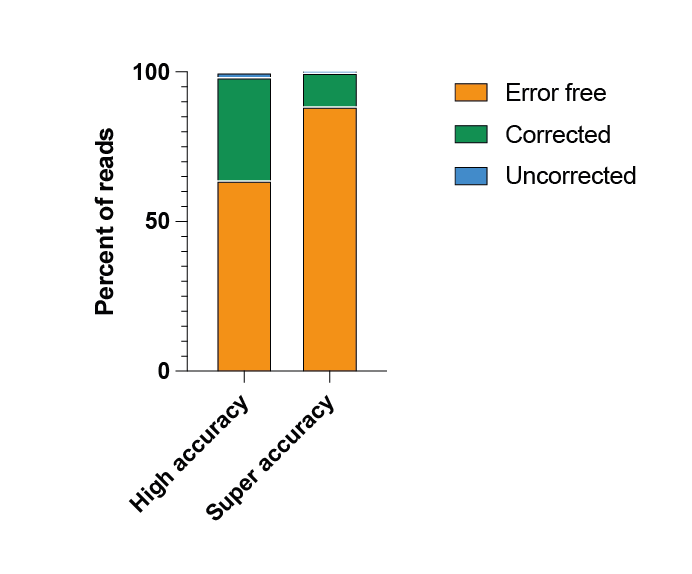

**Figure 12: Improved basecalling accuracy using super accuracy guppy basecalling for ONT technology.**

Percent of CMIs that are correctly sequenced and then error corrected using homotrimer correction using either high accuracy guppy basecalling or super accuracy basecalling for the LSK114 chemistry and R10.4 flow cells.

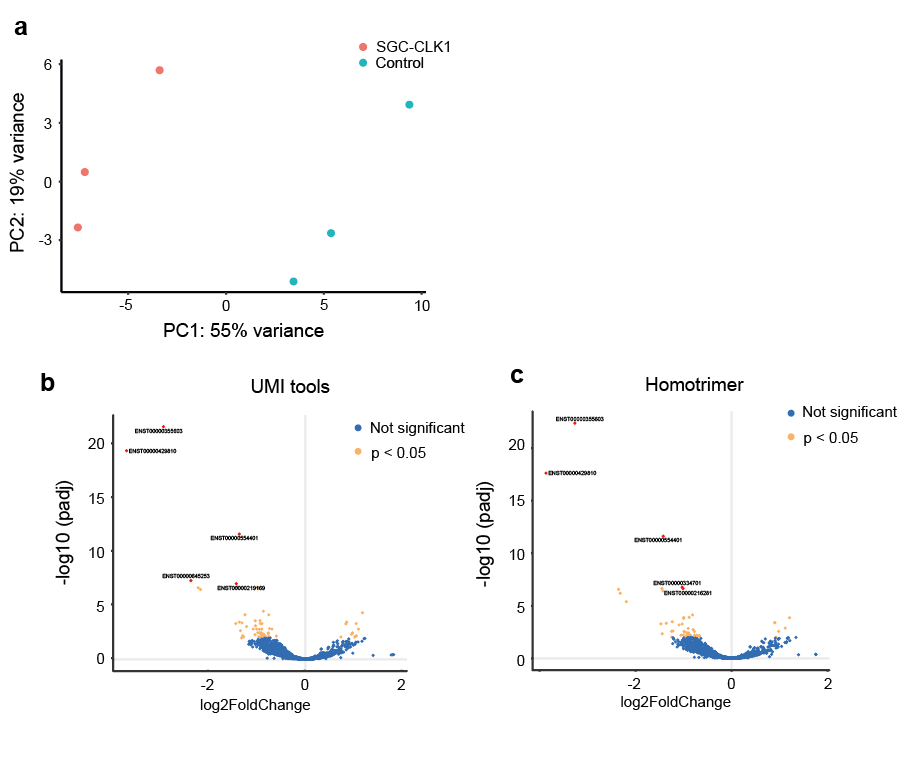

**Figure 13: Analysis of differential gene expression in RM82 Ewing’s sarcoma cells treated with DMSO and CLK-1 inhibitor and sequenced using the ONT platform.**

**a**, A PCR plot showing the variance for cells treated with either DMSO or CLK-1 inhibitor. **b**, A volcano plot showing the log2 fold change and -log10 padj values for cells treated with DMSO or CLK-1 inhibitor, analysed without the inclusion of a UMI during analysis. **d**, A volcano plot showing the log2 fold change and -log10 padj values for cells treated with DMSO or CLK-1 inhibitor and analysed using the homotrimer corrected UMI. These results demonstrate the utility of homotrimer correction in identifying differentially expressed genes and removal of false positive transcripts.

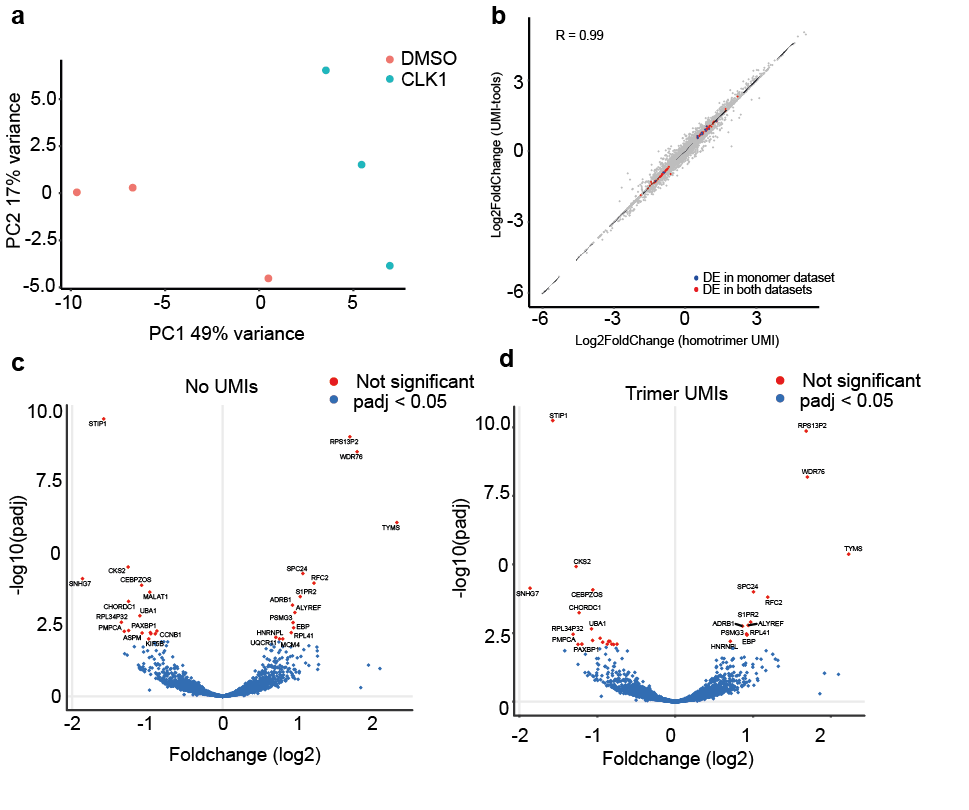

**Figure 14: Analysis of differential gene expression in RM82 Ewing’s sarcoma cells treated with DMSO and CLK-1 inhibitor and sequenced using the Illumina platform.**

**a**, A PCR plot showing the variance for cells treated with either DMSO or CLK-1 inhibitor. **b**, This scatter plot compares the log2 fold changes obtained from randomly collapsing each sequenced trimer UMI with those obtained from homotrimer UMI correction. **c**, A volcano plot showing the log2 fold change and -log10 padj values for cells treated with DMSO or CLK-1 inhibitor, analysed without the inclusion of a UMI during analysis. **d**, A volcano plot showing the log2 fold change and -log10 padj values for cells treated with DMSO or CLK-1 inhibitor and analysed using the homotrimer corrected UMI. These results demonstrate the utility of homotrimer correction in identifying differentially expressed genes and removal of false positive genes.

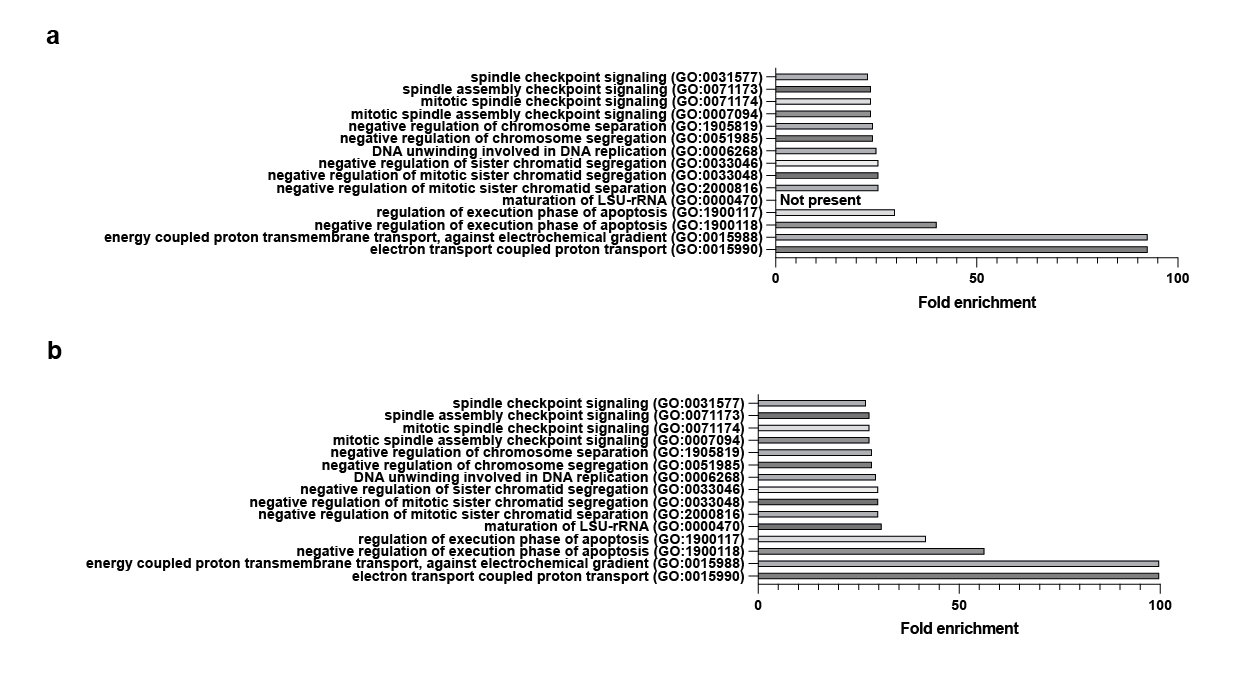

**Figure 15: Go analysis of the differentially expressed genes between DMSO and CLK-1 inhibitor.**

Go analysis was performed for data that was not corrected using UMIs (**a**) and differentially regulated genes following homotrimer correction (**b**).

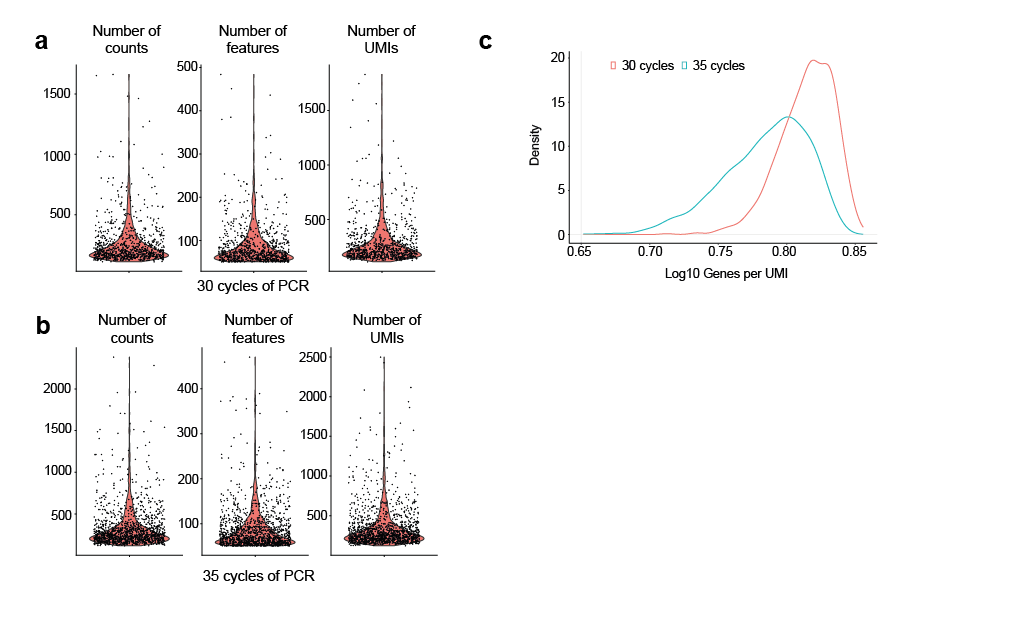

**Figure 16: Quality metrics for 10X chromium single-cell sequencing libraries amplified using 30 and 35 cycles of PCR.**

The number of counts, features and number of UMIs for 10X Chromium libraries PCR amplified for 30 cycles (**a**) and 35 cycles (**b**). Each dot represents a single-cell following filtering. **c**, The log10 genes per UMI plotted as a density.

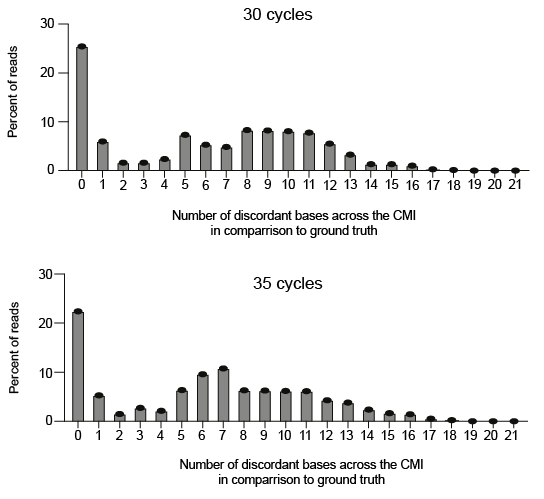

**Figure 17: The number of discordant bases across the full length of the homotrimer UMI**

The number of errors per read between the sequenced CMI and the ground truth CMI following 30 and 35 PCR cycles.

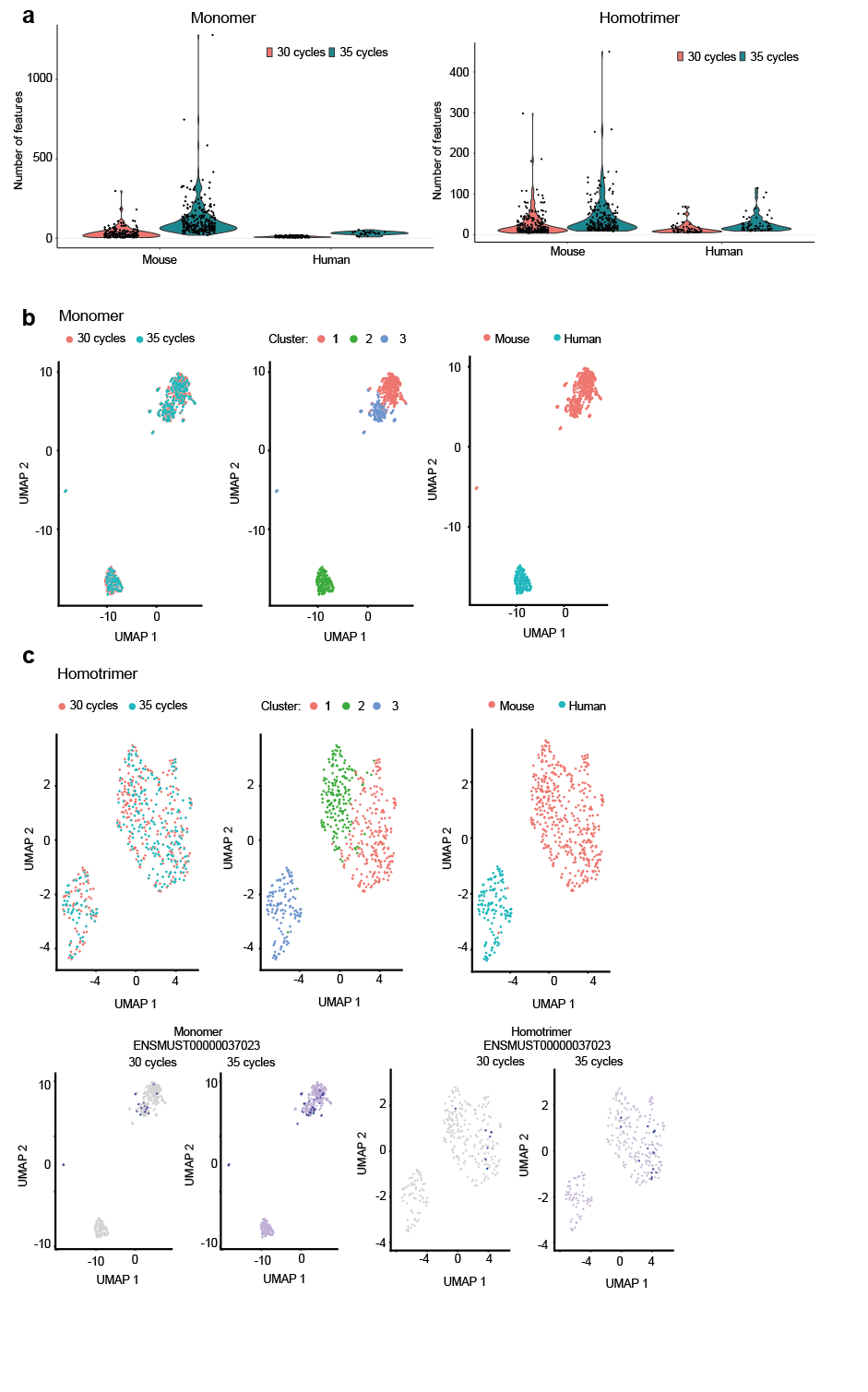

**Figure 18: Number of features and UMAP clustering for the integrated analysis of homotrimer drop-seq UMIs following 30 and 35 cycles of PCR.**

**a**, The number of features detected within the mouse and human cells for monomer (left panel) and homotrimer (right panel) UMIs following 30 and 35 PCR cycles. UMAP plots showing the integration, clustering and annotation of libraries amplified following 30 and 35 PCR cycles for monomer (**b**) and homotrimer (**c**) UMIs. **d**, UMAP plots showing the expression of a non-significant gene ENSMUST0000037023 in monomers (left panels) and homotrimer corrected (right panels) following 30 and 35 cycles of PCR. Even though this gene is not considered significantly different between 30 and 35 cycles in both the monomer and homotrimer datasets, there is generally an overall increase in background counts following 35 cycles of PCR in the monomer dataset that is not apparent within the homotrimer dataset.
