## Supplementary Tables for "Correcting PCR amplification errors in unique molecular identifiers to generate absolute numbers of sequencing molecules"

**Table 1: Barcoding homotrimer primers**

| Barcode | Sequence |
| --- | --- |
| 1 | **AAATTTGGGCCC**AAGCTGTGGTATCAACGCAGAGT |
| 2 | **TTTCCCAAAGGG**AAGCTGTGGTATCAACGCAGAGT |
| 3 | **GGGAAACCCTTT**AAGCTGTGGTATCAACGCAGAGT |
| 4 | **CCCGGGTTTAAA**AAGCTGTGGTATCAACGCAGAGT |

**Table 2: Differentially regulated human genes from 10X chromium experiment.**

|  | p_val | avg_log2FC | pct.1 | pct.2 | p_val_adj |
| --- | --- | --- | --- | --- | --- |
| ENST00000531372.1 | 1.46E-84 | -1.1663112 | 0.876 | 0.989 | 1.33E-80 |
| ENST00000390321.2 | 1.86E-82 | -1.1181074 | 0.91 | 0.994 | 1.70E-78 |
| ENST00000532223.2 | 1.35E-79 | -1.0413469 | 0.946 | 0.995 | 1.23E-75 |
| ENST00000526893.6 | 1.73E-79 | -1.0418797 | 0.946 | 0.995 | 1.58E-75 |
| ENST00000390312.2 | 3.64E-79 | -1.0761879 | 0.925 | 0.992 | 3.32E-75 |
| ENST00000408957.6 | 1.94E-17 | 1.72746934 | 0.37 | 0.16 | 1.77E-13 |
| ENST00000552951.7 | 2.16E-17 | 1.32036377 | 0.516 | 0.33 | 1.97E-13 |
| ENST00000620157.4 | 1.25E-15 | 1.9764611 | 0.304 | 0.12 | 1.14E-11 |
| ENST00000534980.7 | 2.50E-15 | 1.96490342 | 0.302 | 0.12 | 2.28E-11 |
| ENST00000378119.9 | 5.35E-15 | 2.34250392 | 0.221 | 0.063 | 4.87E-11 |
| ENST00000620395.2 | 1.02E-13 | -1.4195983 | 0.075 | 0.265 | 9.27E-10 |
| ENST00000309733.6 | 1.63E-13 | 1.49231239 | 0.406 | 0.233 | 1.48E-09 |
| ENST00000425611.7 | 7.52E-11 | 1.79685504 | 0.226 | 0.089 | 6.85E-07 |
| ENST00000244020.5 | 2.44E-10 | 2.09214051 | 0.158 | 0.045 | 2.22E-06 |
| ENST00000538183.7 | 4.57E-09 | 1.22531603 | 0.382 | 0.251 | 4.16E-05 |
| ENST00000253039.9 | 1.11E-08 | 2.020607 | 0.158 | 0.055 | 0.00010068 |
| ENST00000489060.1 | 1.52E-08 | 1.44334711 | 0.304 | 0.178 | 0.00013809 |
| ENST00000264993.7 | 1.07E-07 | 2.0894513 | 0.144 | 0.052 | 0.00097872 |
| ENST00000430069.5 | 1.53E-07 | 2.1663971 | 0.112 | 0.032 | 0.0013963 |
| ENST00000629042.2 | 1.93E-07 | 1.99979306 | 0.109 | 0.031 | 0.00175802 |
| ENST00000558939.5 | 2.84E-07 | 2.09230461 | 0.102 | 0.028 | 0.00258608 |
| ENST00000616917.4 | 4.64E-07 | 1.86824809 | 0.114 | 0.036 | 0.00422226 |
| ENST00000627388.2 | 5.44E-07 | 2.08500244 | 0.1 | 0.028 | 0.00495735 |
| ENST00000258149.10 | 6.19E-07 | 1.65897531 | 0.175 | 0.079 | 0.00563924 |
| ENST00000262160.11 | 7.77E-07 | 1.78240307 | 0.185 | 0.089 | 0.00707443 |
| ENST00000420115.6 | 1.36E-06 | 1.92090761 | 0.131 | 0.05 | 0.01234361 |
| ENST00000464611.1 | 1.88E-06 | 2.1211892 | 0.112 | 0.039 | 0.01715786 |
| ENST00000633335.1 | 3.52E-06 | 1.25855476 | 0.251 | 0.149 | 0.03206667 |
| ENST00000406053.5 | 4.45E-06 | 1.60091322 | 0.156 | 0.071 | 0.04049086 |
| ENST00000309955.8 | 5.12E-06 | 1.34924064 | 0.178 | 0.087 | 0.0466181 |
| ENST00000254801.9 | 5.43E-06 | 1.22454708 | 0.268 | 0.17 | 0.04943999 |

**Table 3: Differentially regulated mouse genes from 10X chromium experiment**

|  | p_val | avg_log2FC | pct.1 | pct.2 | p_val_adj |
| --- | --- | --- | --- | --- | --- |
| ENSMUST00000019649.4 | 2.84E-49 | -1.1254035 | 0.81 | 0.97 | 2.58E-45 |
| ENSMUST00000034966.9 | 1.29E-23 | -1.5416177 | 0.241 | 0.592 | 1.17E-19 |
| ENSMUST00000027495.15 | 4.64E-15 | 2.14042829 | 0.275 | 0.082 | 4.22E-11 |
| ENSMUST00000179353.8 | 4.83E-15 | 2.13328454 | 0.275 | 0.082 | 4.40E-11 |
| ENSMUST00000168776.8 | 4.94E-15 | 2.13566972 | 0.275 | 0.082 | 4.50E-11 |
| ENSMUST00000172165.8 | 1.11E-14 | 2.09064812 | 0.275 | 0.083 | 1.01E-10 |
| ENSMUST00000190226.7 | 4.62E-14 | 2.32149094 | 0.241 | 0.066 | 4.21E-10 |
| ENSMUST00000035484.11 | 1.65E-13 | 2.18959266 | 0.253 | 0.078 | 1.50E-09 |
| ENSMUST00000072249.13 | 1.93E-13 | 1.97398238 | 0.288 | 0.102 | 1.76E-09 |
| ENSMUST00000116517.9 | 5.06E-13 | 1.98857307 | 0.278 | 0.099 | 4.61E-09 |
| ENSMUST00000244431.1 | 2.07E-11 | -1.6646146 | 0.089 | 0.283 | 1.88E-07 |
| ENSMUST00000198180.5 | 2.93E-11 | 1.76843002 | 0.256 | 0.093 | 2.66E-07 |
| ENSMUST00000197488.5 | 1.29E-10 | 1.82742073 | 0.228 | 0.078 | 1.17E-06 |
| ENSMUST00000029446.13 | 1.63E-09 | 1.66084648 | 0.259 | 0.112 | 1.48E-05 |
| ENSMUST00000103525.3 | 1.87E-08 | -1.0394765 | 0.158 | 0.342 | 0.00017058 |
| ENSMUST00000030578.14 | 2.20E-08 | 1.08845047 | 0.399 | 0.243 | 0.00020039 |
| ENSMUST00000227415.2 | 9.13E-08 | -2.3399698 | 0.025 | 0.135 | 0.00083114 |
| ENSMUST00000165853.2 | 2.07E-07 | 1.05376061 | 0.383 | 0.243 | 0.00188713 |
| ENSMUST00000080726.6 | 9.38E-07 | -1.1822651 | 0.079 | 0.211 | 0.00853962 |
| ENSMUST00000092404.13 | 3.87E-06 | 1.71401331 | 0.108 | 0.03 | 0.03522523 |

**Table 4: Differentially regulated human genes between 30 and 35 cycles following monomer UMI demultiplexing.**

**­­**

|  | p_val | avg_diff | pct.1 | pct.2 | p_val_adj |
| --- | --- | --- | --- | --- | --- |
| ENSMUST00000023599.13 | 2.53E-33 | -0.0909149 | 0.01 | 0.673 | 4.40E-30 |
| ENSMUST00000043864.9 | 3.08E-30 | -0.0347918 | 0.01 | 0.158 | 5.37E-27 |
| ENST00000551018.5 | 3.75E-28 | -0.0579192 | 0.04 | 0.614 | 6.52E-25 |
| ENST00000334478.9 | 3.85E-28 | -0.0558209 | 0.04 | 0.614 | 6.71E-25 |
| ENST00000435275.5 | 5.96E-28 | -0.2129332 | 0.05 | 0.673 | 1.04E-24 |
| ENST00000355731.8 | 5.48E-27 | -0.1155254 | 0.059 | 0.693 | 9.55E-24 |
| ENST00000399075.6 | 1.31E-26 | -0.2413625 | 0.02 | 0.752 | 2.28E-23 |
| ENST00000504110.2 | 1.31E-26 | -0.2413625 | 0.02 | 0.752 | 2.28E-23 |
| ENST00000613865.5 | 1.15E-25 | -0.2222092 | 0.05 | 0.644 | 2.00E-22 |
| ENSMUST00000136342.9 | 5.08E-25 | -0.0485461 | 0.01 | 0.287 | 8.84E-22 |
| ENSMUST00000030613.11 | 5.08E-25 | -0.0543999 | 0.01 | 0.287 | 8.84E-22 |
| ENST00000330494.12 | 5.08E-25 | -0.2032588 | 0.01 | 0.752 | 8.84E-22 |
| ENST00000358181.8 | 5.08E-25 | -0.2032588 | 0.01 | 0.752 | 8.84E-22 |
| ENST00000584577.5 | 1.12E-24 | -0.2511252 | 0.059 | 0.594 | 1.95E-21 |
| ENST00000340913.10 | 1.17E-24 | -0.2304482 | 0.03 | 0.683 | 2.04E-21 |
| ENST00000546500.5 | 1.17E-24 | -0.2304473 | 0.03 | 0.683 | 2.04E-21 |
| ENST00000372360.8 | 1.46E-24 | -0.2246219 | 0.05 | 0.584 | 2.55E-21 |
| ENSMUST00000084846.12 | 8.66E-24 | -0.0463773 | 0.01 | 0.307 | 1.51E-20 |
| ENST00000261479.9 | 2.65E-23 | -0.2104473 | 0.03 | 0.535 | 4.62E-20 |
| ENST00000555764.5 | 2.65E-23 | -0.2104474 | 0.03 | 0.535 | 4.62E-20 |
| ENST00000330752.12 | 2.74E-23 | -0.1843082 | 0.02 | 0.584 | 4.77E-20 |
| ENST00000405805.5 | 5.38E-23 | -0.2878289 | 0.04 | 0.733 | 9.37E-20 |
| ENST00000435983.5 | 1.60E-22 | -0.2297528 | 0.089 | 0.733 | 2.78E-19 |
| ENST00000273223.10 | 1.72E-22 | -0.2191 | 0.089 | 0.733 | 2.99E-19 |
| ENST00000396957.5 | 1.81E-22 | -0.2153185 | 0.089 | 0.752 | 3.15E-19 |
| ENST00000616928.3 | 3.25E-22 | -0.0735431 | 0.059 | 0.693 | 5.66E-19 |
| ENST00000610953.3 | 4.51E-22 | -0.2860542 | 0.069 | 0.822 | 7.85E-19 |
| ENST00000456586.5 | 1.15E-21 | -0.2835875 | 0.069 | 0.752 | 2.00E-18 |
| ENST00000532917.2 | 2.52E-21 | -0.0173951 | 0.03 | 0.406 | 4.39E-18 |
| ENST00000471855.1 | 3.33E-21 | -0.0479422 | 0.059 | 0.644 | 5.79E-18 |
| ENST00000449470.6 | 4.42E-21 | -0.0832719 | 0.05 | 0.535 | 7.70E-18 |
| ENST00000357865.6 | 9.59E-21 | -0.1237055 | 0.069 | 0.505 | 1.67E-17 |
| ENST00000302182.8 | 1.07E-20 | 0.04958333 | 0.069 | 0.356 | 1.86E-17 |
| ENST00000395837.1 | 1.30E-20 | 0.04135911 | 0.059 | 0.475 | 2.26E-17 |
| ENST00000614404.1 | 1.30E-20 | 0.04553576 | 0.059 | 0.475 | 2.26E-17 |
| ENST00000455785.7 | 1.48E-20 | -0.1540661 | 0.089 | 0.703 | 2.58E-17 |
| ENST00000399728.5 | 1.52E-20 | -0.1537927 | 0.089 | 0.703 | 2.65E-17 |
| ENST00000225655.5 | 2.45E-20 | -0.2255804 | 0.109 | 0.663 | 4.27E-17 |
| ENST00000274242.10 | 3.10E-20 | -0.1176062 | 0.03 | 0.545 | 5.40E-17 |
| ENST00000395839.5 | 3.36E-20 | 0.01745546 | 0.05 | 0.366 | 5.84E-17 |
| ENST00000645496.1 | 4.33E-20 | -0.0411846 | 0.05 | 0.554 | 7.54E-17 |
| ENST00000393394.5 | 4.43E-20 | -0.0865175 | 0.05 | 0.545 | 7.72E-17 |
| ENST00000646648.1 | 6.10E-20 | -0.2109676 | 0.01 | 0.762 | 1.06E-16 |
| ENST00000533773.5 | 6.10E-20 | -0.2109684 | 0.01 | 0.762 | 1.06E-16 |
| ENST00000323688.10 | 6.10E-20 | -0.2109499 | 0.01 | 0.762 | 1.06E-16 |
| ENST00000529010.6 | 6.10E-20 | -0.2109648 | 0.01 | 0.762 | 1.06E-16 |
| ENST00000512737.5 | 6.84E-20 | -0.1616641 | 0.03 | 0.624 | 1.19E-16 |
| ENST00000503064.5 | 6.84E-20 | -0.1616648 | 0.03 | 0.624 | 1.19E-16 |
| ENST00000512088.1 | 7.04E-20 | -0.2262655 | 0.01 | 0.693 | 1.23E-16 |
| ENST00000252102.8 | 7.04E-20 | -0.226266 | 0.01 | 0.693 | 1.23E-16 |
| ENST00000502960.1 | 7.04E-20 | -0.226266 | 0.01 | 0.693 | 1.23E-16 |
| ENST00000359681.3 | 1.07E-19 | -0.1614745 | 0.079 | 0.495 | 1.85E-16 |
| ENST00000491306.6 | 1.09E-19 | -0.166695 | 0.079 | 0.495 | 1.90E-16 |
| ENST00000558131.1 | 1.77E-19 | -0.1464067 | 0.04 | 0.594 | 3.09E-16 |
| ENST00000444681.6 | 1.91E-19 | -0.0939576 | 0.03 | 0.782 | 3.32E-16 |
| ENST00000302907.9 | 2.70E-19 | -0.2522946 | 0.079 | 0.812 | 4.71E-16 |
| ENST00000616839.3 | 2.83E-19 | -0.1893624 | 0.079 | 0.822 | 4.93E-16 |
| ENST00000084795.9 | 3.92E-19 | -0.1611391 | 0.059 | 0.525 | 6.83E-16 |
| ENST00000295955.13 | 4.09E-19 | -0.0924181 | 0.05 | 0.545 | 7.12E-16 |
| ENST00000559082.5 | 4.69E-19 | -0.0783901 | 0.04 | 0.535 | 8.16E-16 |
| ENST00000370321.8 | 7.65E-19 | -0.0740323 | 0.03 | 0.594 | 1.33E-15 |
| ENST00000596046.1 | 8.49E-19 | -0.1877205 | 0.119 | 0.634 | 1.48E-15 |
| ENST00000429711.7 | 1.11E-18 | -0.2170964 | 0.089 | 0.624 | 1.93E-15 |
| ENST00000367679.7 | 1.24E-18 | -0.1102098 | 0.059 | 0.713 | 2.16E-15 |
| ENST00000368167.8 | 1.65E-18 | -0.0258499 | 0.04 | 0.673 | 2.88E-15 |
| ENST00000360830.9 | 2.88E-18 | -0.1999409 | 0.03 | 0.475 | 5.01E-15 |
| ENSMUST00000103418.3 | 3.04E-18 | -0.006052 | 0.05 | 0.109 | 5.30E-15 |
| ENST00000648640.1 | 6.50E-18 | -0.1766335 | 0.079 | 0.525 | 1.13E-14 |
| ENST00000576917.5 | 8.27E-18 | -0.1388421 | 0.03 | 0.594 | 1.44E-14 |
| ENST00000529920.5 | 1.06E-17 | -0.1765355 | 0.069 | 0.584 | 1.84E-14 |
| ENST00000342192.8 | 1.08E-17 | -0.2111848 | 0.04 | 0.406 | 1.88E-14 |
| ENST00000458578.6 | 1.08E-17 | -0.1559374 | 0.04 | 0.436 | 1.88E-14 |
| ENST00000496722.1 | 1.08E-17 | -0.1559374 | 0.04 | 0.436 | 1.88E-14 |
| ENST00000376236.9 | 1.15E-17 | -0.1226653 | 0.02 | 0.574 | 2.00E-14 |
| ENST00000303127.12 | 1.26E-17 | -0.0405958 | 0.04 | 0.564 | 2.19E-14 |
| ENST00000549920.6 | 2.79E-17 | -0.1622408 | 0.059 | 0.525 | 4.85E-14 |
| ENST00000549370.5 | 2.79E-17 | -0.1629762 | 0.059 | 0.525 | 4.85E-14 |
| ENST00000575842.5 | 3.49E-17 | -0.0001119 | 0.04 | 0.673 | 6.07E-14 |
| ENST00000230050.4 | 4.15E-17 | -0.2448091 | 0.109 | 0.564 | 7.22E-14 |
| ENSMUST00000190686.7 | 4.38E-17 | -0.0799728 | 0.01 | 0.396 | 7.62E-14 |
| ENST00000526248.5 | 5.42E-17 | -0.243731 | 0.129 | 0.822 | 9.44E-14 |
| ENST00000338639.10 | 5.55E-17 | -0.1514323 | 0.02 | 0.495 | 9.66E-14 |
| ENST00000617605.4 | 8.71E-17 | -0.1225784 | 0.059 | 0.564 | 1.52E-13 |
| ENST00000311111.11 | 1.27E-16 | -0.2909645 | 0.05 | 0.485 | 2.21E-13 |
| ENST00000404735.1 | 1.36E-16 | 0.19733387 | 0.05 | 0.149 | 2.36E-13 |
| ENST00000480306.5 | 1.39E-16 | -0.2512742 | 0.069 | 0.564 | 2.42E-13 |
| ENST00000290299.7 | 1.62E-16 | -0.1948099 | 0.04 | 0.663 | 2.81E-13 |
| ENST00000652380.1 | 1.62E-16 | -0.1948099 | 0.04 | 0.663 | 2.81E-13 |
| ENST00000497035.5 | 1.67E-16 | -0.1491818 | 0.079 | 0.495 | 2.90E-13 |
| ENST00000454893.1 | 2.88E-16 | -0.1283881 | 0.04 | 0.663 | 5.01E-13 |
| ENST00000528957.5 | 4.50E-16 | -0.2440061 | 0.079 | 0.644 | 7.84E-13 |
| ENST00000262584.7 | 4.79E-16 | -0.2441851 | 0.079 | 0.644 | 8.35E-13 |
| ENST00000394920.6 | 4.89E-16 | -0.2443749 | 0.079 | 0.644 | 8.52E-13 |
| ENST00000493678.5 | 4.95E-16 | -0.1488518 | 0.02 | 0.485 | 8.61E-13 |
| ENST00000341423.10 | 6.40E-16 | -0.2766551 | 0.109 | 0.693 | 1.11E-12 |
| ENST00000616877.3 | 1.48E-15 | -0.1448874 | 0.059 | 0.693 | 2.58E-12 |
| ENST00000485390.5 | 1.49E-15 | -0.1396368 | 0.129 | 0.475 | 2.60E-12 |
| ENST00000515770.2 | 2.17E-15 | 0.1378334 | 0.04 | 0.178 | 3.77E-12 |
| ENST00000652689.1 | 2.28E-15 | 0.13608385 | 0.04 | 0.178 | 3.97E-12 |
| ENST00000321153.9 | 2.65E-15 | -0.269015 | 0.04 | 0.297 | 4.61E-12 |
| ENST00000294189.11 | 3.32E-15 | -0.1877578 | 0.079 | 0.564 | 5.77E-12 |
| ENST00000380358.9 | 6.41E-15 | -0.1381672 | 0.01 | 0.673 | 1.12E-11 |
| ENST00000380842.5 | 7.74E-15 | 0.14153102 | 0.02 | 0.158 | 1.35E-11 |
| ENST00000460403.1 | 7.74E-15 | 0.14153102 | 0.02 | 0.158 | 1.35E-11 |
| ENST00000513345.5 | 1.36E-14 | -0.0833208 | 0.04 | 0.673 | 2.36E-11 |
| ENST00000507591.2 | 1.60E-14 | 0.12409084 | 0.04 | 0.178 | 2.78E-11 |
| ENST00000539281.5 | 2.47E-14 | -0.2565793 | 0.01 | 0.703 | 4.29E-11 |
| ENST00000326004.4 | 4.50E-14 | -0.2759627 | 0.069 | 0.634 | 7.84E-11 |
| ENST00000620720.3 | 5.05E-14 | -0.072474 | 0.069 | 0.634 | 8.80E-11 |
| ENST00000587367.5 | 5.60E-14 | -0.0696452 | 0.03 | 0.713 | 9.75E-11 |
| ENST00000342669.7 | 5.60E-14 | -0.0696481 | 0.03 | 0.713 | 9.75E-11 |
| ENST00000588301.5 | 5.60E-14 | -0.0696518 | 0.03 | 0.713 | 9.75E-11 |
| ENST00000672405.1 | 5.73E-14 | -0.225207 | 0.03 | 0.624 | 9.98E-11 |
| ENST00000422447.8 | 5.73E-14 | -0.2261397 | 0.03 | 0.624 | 9.98E-11 |
| ENST00000550920.6 | 5.86E-14 | 0.05587003 | 0.01 | 0.188 | 1.02E-10 |
| ENST00000370089.6 | 5.96E-14 | -0.1562345 | 0.05 | 0.356 | 1.04E-10 |
| ENST00000396856.5 | 6.15E-14 | -0.2887148 | 0.069 | 0.624 | 1.07E-10 |
| ENST00000420712.1 | 6.16E-14 | -0.2205914 | 0.059 | 0.416 | 1.07E-10 |
| ENST00000509698.5 | 6.19E-14 | -0.1506441 | 0.05 | 0.366 | 1.08E-10 |
| ENST00000370081.6 | 6.19E-14 | -0.1506514 | 0.05 | 0.366 | 1.08E-10 |
| ENST00000419376.2 | 7.08E-14 | -0.2032683 | 0.01 | 0.673 | 1.23E-10 |
| ENST00000361575.4 | 8.63E-14 | -0.2950988 | 0.079 | 0.723 | 1.50E-10 |
| ENSMUST00000099047.4 | 1.00E-13 | -0.0849234 | 0.01 | 0.356 | 1.74E-10 |
| ENST00000351839.7 | 1.59E-13 | 0.2235644 | 1 | 0.525 | 2.76E-10 |
| ENST00000360384.9 | 1.59E-13 | 0.22356434 | 1 | 0.525 | 2.76E-10 |
| ENST00000558397.1 | 1.91E-13 | -0.1918567 | 0.099 | 0.545 | 3.32E-10 |
| ENST00000589037.5 | 1.95E-13 | -0.3215923 | 0.059 | 0.693 | 3.39E-10 |
| ENST00000579226.5 | 2.10E-13 | -0.0915637 | 0.02 | 0.574 | 3.65E-10 |
| ENST00000304414.12 | 2.53E-13 | 0.00657722 | 0.04 | 0.545 | 4.41E-10 |
| ENST00000229270.8 | 2.59E-13 | -0.1121618 | 0.04 | 0.307 | 4.51E-10 |
| ENST00000613953.4 | 2.59E-13 | -0.1121618 | 0.04 | 0.307 | 4.51E-10 |
| ENST00000396705.10 | 2.62E-13 | -0.1097842 | 0.04 | 0.307 | 4.56E-10 |
| ENST00000396651.8 | 2.87E-13 | -0.1423853 | 0.119 | 0.396 | 5.00E-10 |
| ENST00000459637.2 | 3.65E-13 | -0.1526593 | 0.05 | 0.347 | 6.35E-10 |
| ENST00000251453.8 | 5.44E-13 | 0.14564861 | 0.03 | 0.139 | 9.47E-10 |
| ENST00000339471.8 | 5.76E-13 | 0.1405812 | 0.03 | 0.139 | 1.00E-09 |
| ENST00000361453.3 | 8.27E-13 | 0.15648838 | 0.03 | 0.228 | 1.44E-09 |
| ENST00000361851.1 | 9.26E-13 | -0.3537809 | 0.059 | 0.673 | 1.61E-09 |
| ENST00000544417.5 | 9.27E-13 | -0.2009801 | 0.069 | 0.653 | 1.61E-09 |
| ENST00000589913.6 | 9.98E-13 | -0.2343406 | 0.069 | 0.683 | 1.74E-09 |
| ENST00000253788.11 | 9.98E-13 | -0.2481026 | 0.069 | 0.683 | 1.74E-09 |
| ENST00000562336.5 | 1.31E-12 | -0.2473453 | 0.05 | 0.772 | 2.28E-09 |
| ENST00000253452.7 | 1.31E-12 | -0.2373276 | 0.05 | 0.762 | 2.28E-09 |
| ENST00000561569.5 | 1.31E-12 | -0.237328 | 0.05 | 0.762 | 2.28E-09 |
| ENST00000617731.2 | 3.22E-12 | -0.145329 | 0.119 | 0.495 | 5.61E-09 |
| ENST00000647841.1 | 3.28E-12 | -0.1444834 | 0.119 | 0.495 | 5.71E-09 |
| ENST00000633731.1 | 3.40E-12 | -0.148528 | 0.119 | 0.505 | 5.92E-09 |
| ENST00000361427.6 | 3.83E-12 | 0.14463452 | 0.079 | 0.198 | 6.67E-09 |
| ENST00000648006.2 | 4.94E-12 | -0.2006889 | 0.069 | 0.653 | 8.60E-09 |
| ENST00000619352.4 | 5.12E-12 | 0.13354895 | 0.069 | 0.208 | 8.91E-09 |
| ENST00000396076.5 | 5.45E-12 | -0.0752274 | 0.04 | 0.594 | 9.49E-09 |
| ENST00000673047.1 | 5.45E-12 | -0.093982 | 0.04 | 0.614 | 9.49E-09 |
| ENST00000350669.5 | 5.45E-12 | -0.0906579 | 0.04 | 0.614 | 9.49E-09 |
| ENST00000479563.5 | 5.50E-12 | -0.1571941 | 0.059 | 0.455 | 9.58E-09 |
| ENST00000409360.1 | 7.16E-12 | -0.1234057 | 0.03 | 0.604 | 1.25E-08 |
| ENST00000369713.10 | 8.20E-12 | -0.2366677 | 0.01 | 0.673 | 1.43E-08 |
| ENST00000489459.5 | 8.33E-12 | -0.1668419 | 0.109 | 0.554 | 1.45E-08 |
| ENST00000578611.5 | 1.14E-11 | -0.0716723 | 0.02 | 0.535 | 1.98E-08 |
| ENST00000484616.2 | 1.17E-11 | -0.2286684 | 0.099 | 0.545 | 2.04E-08 |
| ENST00000530945.1 | 1.24E-11 | 0.37248732 | 0.03 | 0.198 | 2.16E-08 |
| ENST00000308162.10 | 1.25E-11 | 0.27730971 | 0.03 | 0.178 | 2.18E-08 |
| ENST00000507754.9 | 1.34E-11 | -0.1049764 | 0.05 | 0.376 | 2.33E-08 |
| ENST00000322723.9 | 1.57E-11 | -0.1621613 | 0.05 | 0.406 | 2.73E-08 |
| ENST00000361681.2 | 1.65E-11 | -0.1931535 | 0.05 | 0.475 | 2.87E-08 |
| ENST00000531188.6 | 1.80E-11 | -0.2249413 | 0.168 | 0.792 | 3.13E-08 |
| ENST00000278572.10 | 1.93E-11 | -0.2409991 | 0.158 | 0.752 | 3.36E-08 |
| ENST00000530721.5 | 1.99E-11 | -0.2164233 | 0.168 | 0.782 | 3.46E-08 |
| ENST00000376049.4 | 2.32E-11 | -0.192463 | 0.01 | 0.673 | 4.04E-08 |
| ENST00000233892.8 | 4.10E-11 | -0.1219059 | 0.03 | 0.604 | 7.13E-08 |
| ENST00000448447.6 | 4.10E-11 | -0.1245424 | 0.03 | 0.604 | 7.13E-08 |
| ENST00000323303.9 | 4.10E-11 | -0.1254758 | 0.03 | 0.604 | 7.13E-08 |
| ENST00000368166.7 | 4.45E-11 | -0.0644246 | 0.01 | 0.624 | 7.75E-08 |
| ENST00000467168.5 | 5.50E-11 | 0.00854313 | 0.158 | 0.248 | 9.58E-08 |
| ENST00000502506.6 | 7.10E-11 | -0.1108549 | 0.05 | 0.376 | 1.24E-07 |
| ENST00000361335.1 | 7.83E-11 | -0.0977323 | 0.089 | 0.525 | 1.36E-07 |
| ENST00000429728.1 | 1.30E-10 | -0.1339823 | 0.03 | 0.584 | 2.26E-07 |
| ENST00000396203.7 | 1.44E-10 | -0.150113 | 0.059 | 0.446 | 2.51E-07 |
| ENST00000619603.1 | 1.65E-10 | -0.1334485 | 0.059 | 0.515 | 2.87E-07 |
| ENST00000525451.6 | 1.67E-10 | 0.29073419 | 0.04 | 0.168 | 2.90E-07 |
| ENST00000468812.6 | 1.81E-10 | -0.1594586 | 0.109 | 0.545 | 3.16E-07 |
| ENST00000355968.10 | 1.81E-10 | -0.1570541 | 0.109 | 0.554 | 3.16E-07 |
| ENST00000540844.5 | 1.84E-10 | -0.1675401 | 0.03 | 0.198 | 3.21E-07 |
| ENST00000470450.5 | 1.91E-10 | -0.1599976 | 0.03 | 0.198 | 3.32E-07 |
| ENST00000286788.9 | 1.91E-10 | -0.1061333 | 0.03 | 0.208 | 3.32E-07 |
| ENST00000598495.5 | 2.91E-10 | -0.1699578 | 0.158 | 0.564 | 5.07E-07 |
| ENST00000601521.5 | 2.91E-10 | -0.1699578 | 0.158 | 0.564 | 5.07E-07 |
| ENST00000196551.8 | 2.91E-10 | -0.1693747 | 0.158 | 0.564 | 5.07E-07 |
| ENST00000626906.1 | 3.09E-10 | 0.12707936 | 0.03 | 0.178 | 5.38E-07 |
| ENST00000465752.1 | 3.09E-10 | 0.12707507 | 0.03 | 0.178 | 5.38E-07 |
| ENST00000379143.10 | 3.82E-10 | -0.2141326 | 0.03 | 0.713 | 6.65E-07 |
| ENST00000379160.3 | 3.82E-10 | -0.2141326 | 0.03 | 0.713 | 6.65E-07 |
| ENST00000645674.2 | 4.06E-10 | -0.2012685 | 0.079 | 0.634 | 7.08E-07 |
| ENST00000600659.3 | 4.27E-10 | -0.0739359 | 0.079 | 0.178 | 7.43E-07 |
| ENST00000217652.7 | 4.47E-10 | -0.0576914 | 0.02 | 0.465 | 7.78E-07 |
| ENST00000369817.6 | 7.46E-10 | 0.02301198 | 0.188 | 0.257 | 1.30E-06 |
| ENST00000322203.7 | 7.88E-10 | 0.04660317 | 0.119 | 0.198 | 1.37E-06 |
| ENST00000356769.7 | 8.81E-10 | 0.08791798 | 0.01 | 0.238 | 1.53E-06 |
| ENST00000381359.5 | 1.13E-09 | -0.2420359 | 0.04 | 0.554 | 1.97E-06 |
| ENST00000249786.8 | 1.13E-09 | -0.2420351 | 0.04 | 0.554 | 1.97E-06 |
| ENST00000457341.6 | 1.33E-09 | -0.1691005 | 0.238 | 0.416 | 2.32E-06 |
| ENST00000332211.10 | 1.75E-09 | -0.0264813 | 0.03 | 0.485 | 3.04E-06 |
| ENST00000403564.5 | 1.76E-09 | -0.1995269 | 0.079 | 0.634 | 3.07E-06 |
| ENST00000646909.1 | 1.82E-09 | -0.1919696 | 0.079 | 0.624 | 3.16E-06 |
| ENST00000462576.5 | 1.82E-09 | -0.2029141 | 0.079 | 0.634 | 3.16E-06 |
| ENST00000394077.8 | 1.88E-09 | -0.1636913 | 0.03 | 0.475 | 3.27E-06 |
| ENST00000464595.1 | 1.91E-09 | -0.1549981 | 0.03 | 0.475 | 3.33E-06 |
| ENST00000579248.5 | 1.94E-09 | -0.0937112 | 0.069 | 0.564 | 3.37E-06 |
| ENST00000275603.9 | 3.00E-09 | -0.1507256 | 0.03 | 0.554 | 5.22E-06 |
| ENST00000338970.10 | 3.11E-09 | -0.1945237 | 0.059 | 0.446 | 5.42E-06 |
| ENST00000614077.4 | 4.03E-09 | -0.0975251 | 0.079 | 0.475 | 7.01E-06 |
| ENST00000481928.1 | 4.52E-09 | -0.1803536 | 0.05 | 0.307 | 7.86E-06 |
| ENST00000479992.5 | 5.43E-09 | -0.2078935 | 0.089 | 0.416 | 9.45E-06 |
| ENST00000600213.3 | 5.93E-09 | -0.1472893 | 0.149 | 0.198 | 1.03E-05 |
| ENST00000331523.6 | 6.10E-09 | 0.00160636 | 0.198 | 0.267 | 1.06E-05 |
| ENST00000633705.1 | 6.52E-09 | 0.01412441 | 0.059 | 0.208 | 1.14E-05 |
| ENST00000632136.1 | 6.52E-09 | 0.01412445 | 0.059 | 0.208 | 1.14E-05 |
| ENST00000580261.5 | 7.92E-09 | -0.1522048 | 0.069 | 0.564 | 1.38E-05 |
| ENST00000370995.6 | 9.76E-09 | -0.0908159 | 0.02 | 0.436 | 1.70E-05 |
| ENST00000518850.5 | 1.16E-08 | -0.2705475 | 0.099 | 0.436 | 2.03E-05 |
| ENST00000489915.1 | 1.43E-08 | -0.2219959 | 0.079 | 0.663 | 2.49E-05 |
| ENST00000219150.9 | 1.62E-08 | 0.20282585 | 0.03 | 0.257 | 2.82E-05 |
| ENST00000630979.3 | 1.64E-08 | -0.1401149 | 0.05 | 0.545 | 2.86E-05 |
| ENST00000323345.11 | 1.64E-08 | -0.1401135 | 0.05 | 0.545 | 2.86E-05 |
| ENST00000625669.2 | 1.64E-08 | -0.1407805 | 0.05 | 0.545 | 2.86E-05 |
| ENST00000463740.5 | 1.64E-08 | -0.1424067 | 0.05 | 0.525 | 2.86E-05 |
| ENST00000272274.8 | 1.99E-08 | 0.11410789 | 0.198 | 0.248 | 3.47E-05 |
| ENST00000599539.5 | 2.06E-08 | 0.0515262 | 0.01 | 0.257 | 3.59E-05 |
| ENST00000422514.7 | 2.90E-08 | -0.2173665 | 0.079 | 0.416 | 5.06E-05 |
| ENST00000424325.6 | 2.96E-08 | 0.02273158 | 0.198 | 0.267 | 5.16E-05 |
| ENST00000233143.6 | 3.08E-08 | -0.0567551 | 0.079 | 0.228 | 5.36E-05 |
| ENST00000287038.8 | 3.22E-08 | -0.2379131 | 0.109 | 0.416 | 5.60E-05 |
| ENST00000554455.5 | 3.55E-08 | -0.04685 | 0.03 | 0.693 | 6.18E-05 |
| ENST00000361891.9 | 3.74E-08 | -0.1585181 | 0.02 | 0.594 | 6.51E-05 |
| ENST00000615950.4 | 3.74E-08 | -0.1585181 | 0.02 | 0.594 | 6.51E-05 |
| ENST00000335503.3 | 4.35E-08 | -0.1096114 | 0.02 | 0.594 | 7.57E-05 |
| ENST00000521291.5 | 4.90E-08 | -0.23513 | 0.099 | 0.426 | 8.53E-05 |
| ENST00000319826.8 | 5.46E-08 | 0.08709229 | 0.198 | 0.267 | 9.50E-05 |
| ENST00000370990.5 | 5.55E-08 | -0.0688449 | 0.02 | 0.426 | 9.66E-05 |
| ENST00000361219.10 | 5.55E-08 | -0.0688484 | 0.02 | 0.426 | 9.66E-05 |
| ENST00000370994.8 | 5.55E-08 | -0.0688484 | 0.02 | 0.426 | 9.66E-05 |
| ENST00000576008.5 | 5.56E-08 | -0.2668138 | 0.129 | 0.505 | 9.68E-05 |
| ENST00000508495.5 | 5.81E-08 | -0.1471348 | 0.059 | 0.624 | 0.00010117 |
| ENST00000477624.1 | 6.13E-08 | -0.114495 | 0.03 | 0.376 | 0.00010681 |
| ENST00000311549.10 | 6.63E-08 | 0.05819579 | 0.198 | 0.277 | 0.00011548 |
| ENST00000553300.5 | 6.77E-08 | -0.0126753 | 0.03 | 0.663 | 0.00011779 |
| ENST00000495383.5 | 9.21E-08 | -0.1878998 | 0.059 | 0.485 | 0.00016027 |
| ENST00000393891.8 | 9.88E-08 | -0.1759333 | 0.059 | 0.505 | 0.00017207 |
| ENST00000647408.1 | 1.03E-07 | -0.043553 | 0.03 | 0.584 | 0.00017852 |
| ENST00000377803.3 | 1.06E-07 | 0.02537817 | 0.139 | 0.228 | 0.00018539 |
| ENST00000538183.7 | 1.20E-07 | -0.1218627 | 0.02 | 0.604 | 0.00020892 |
| ENST00000309268.11 | 1.64E-07 | -0.0074349 | 0.218 | 0.277 | 0.00028574 |
| ENST00000616577.4 | 1.88E-07 | -0.1497251 | 0.04 | 0.426 | 0.00032654 |
| ENST00000649899.1 | 2.28E-07 | 0.01140191 | 0.02 | 0.604 | 0.00039779 |
| ENST00000626972.2 | 2.29E-07 | -0.2141097 | 0.02 | 0.158 | 0.00039863 |
| ENST00000506126.5 | 2.51E-07 | -0.0356909 | 0.149 | 0.208 | 0.00043659 |
| ENST00000276062.8 | 3.01E-07 | -0.1776452 | 0.089 | 0.683 | 0.00052452 |
| ENST00000227378.7 | 3.02E-07 | -0.008203 | 0.05 | 0.614 | 0.00052499 |
| ENST00000552540.5 | 3.84E-07 | -0.0178656 | 0.059 | 0.198 | 0.0006689 |
| ENST00000610020.2 | 4.09E-07 | -0.1674359 | 0.01 | 0.426 | 0.00071143 |
| ENST00000399494.5 | 5.14E-07 | -0.1651387 | 0.03 | 0.436 | 0.00089413 |
| ENST00000318607.10 | 5.27E-07 | -0.1411128 | 0.069 | 0.604 | 0.00091778 |
| ENST00000202773.13 | 7.18E-07 | -0.1873676 | 0.079 | 0.475 | 0.00124958 |
| ENST00000424576.6 | 7.18E-07 | -0.1873681 | 0.079 | 0.475 | 0.00124958 |
| ENST00000553110.7 | 7.46E-07 | -0.1027594 | 0.079 | 0.475 | 0.00129937 |
| ENST00000429437.5 | 9.11E-07 | -0.1060739 | 0.04 | 0.465 | 0.00158614 |
| ENST00000376263.8 | 9.62E-07 | 0.22382962 | 0.01 | 0.257 | 0.00167502 |
| ENST00000376281.8 | 9.62E-07 | 0.22384267 | 0.01 | 0.257 | 0.00167502 |
| ENST00000377811.3 | 9.96E-07 | -0.1776781 | 0.089 | 0.683 | 0.00173391 |
| ENST00000534624.6 | 1.07E-06 | -0.0320549 | 0.05 | 0.604 | 0.00186686 |
| ENST00000527871.5 | 1.08E-06 | -0.1820498 | 0.05 | 0.208 | 0.00187895 |
| ENST00000344746.8 | 1.24E-06 | 0.02210564 | 0.208 | 0.267 | 0.00215285 |
| ENST00000479035.7 | 1.51E-06 | -0.156385 | 0.05 | 0.386 | 0.00262255 |
| ENST00000245857.9 | 1.51E-06 | -0.1600854 | 0.05 | 0.386 | 0.00262255 |
| ENST00000316292.13 | 1.55E-06 | -0.0039712 | 0.208 | 0.257 | 0.00269571 |
| ENST00000514057.1 | 1.75E-06 | -0.3181943 | 0.069 | 0.396 | 0.00305086 |
| ENST00000261700.8 | 1.86E-06 | -0.0291373 | 0.05 | 0.525 | 0.00323187 |
| ENST00000448864.6 | 2.36E-06 | -0.1060957 | 0.059 | 0.426 | 0.00410458 |
| ENST00000647248.2 | 2.36E-06 | -0.1110934 | 0.059 | 0.416 | 0.00410458 |
| ENST00000477151.2 | 2.48E-06 | -0.0132897 | 0.059 | 0.218 | 0.00431424 |
| ENST00000618619.4 | 2.74E-06 | -0.1633372 | 0.05 | 0.545 | 0.00477339 |
| ENST00000461690.5 | 2.79E-06 | 0.07289323 | 0.139 | 0.208 | 0.00486384 |
| ENST00000371554.2 | 3.05E-06 | -0.0710135 | 0.089 | 0.168 | 0.00531319 |
| ENST00000415933.5 | 3.25E-06 | -0.2029361 | 0.089 | 0.535 | 0.00564979 |
| ENST00000381140.9 | 3.98E-06 | -0.0747825 | 0.02 | 0.406 | 0.0069294 |
| ENST00000615060.4 | 4.09E-06 | -0.2437186 | 0.119 | 0.455 | 0.00712343 |
| ENST00000444376.7 | 4.34E-06 | 0.24071624 | 0.02 | 0.307 | 0.00755345 |
| ENST00000497825.5 | 4.59E-06 | -0.0503961 | 0.05 | 0.119 | 0.00799535 |
| ENST00000395119.7 | 7.59E-06 | 0.03073408 | 0.03 | 0.337 | 0.01321111 |
| ENST00000529272.5 | 7.59E-06 | 0.02811707 | 0.03 | 0.337 | 0.01321111 |
| ENST00000632587.1 | 7.59E-06 | 0.02811707 | 0.03 | 0.337 | 0.01321111 |
| ENST00000614575.4 | 7.59E-06 | 0.0281203 | 0.03 | 0.337 | 0.01321111 |
| ENST00000573283.7 | 7.93E-06 | -0.0652091 | 0.04 | 0.545 | 0.01380901 |
| ENST00000366839.8 | 1.21E-05 | -0.0269607 | 0.02 | 0.158 | 0.0209977 |
| ENST00000651761.1 | 1.25E-05 | 0.06496279 | 0.02 | 0.129 | 0.02182321 |
| ENST00000394938.8 | 1.57E-05 | -0.2201783 | 0.069 | 0.396 | 0.02740416 |
| ENST00000592588.6 | 1.66E-05 | -0.1834288 | 0.198 | 0.564 | 0.02898127 |
| ENST00000367142.5 | 1.82E-05 | -0.1857392 | 0.05 | 0.178 | 0.03169516 |
| ENST00000228140.6 | 1.83E-05 | 0.04804688 | 0.069 | 0.208 | 0.03180682 |
| ENST00000569365.6 | 2.17E-05 | -0.1425635 | 0.158 | 0.426 | 0.03783777 |
| ENST00000646664.1 | 2.19E-05 | -0.1895698 | 0.059 | 0.594 | 0.03814112 |
